# Neuronal-class specific molecular cues drive differential myelination in the neocortex

**DOI:** 10.1101/2024.02.20.581268

**Authors:** Vahbiz Jokhi, Nuria Domínguez-Iturza, Kwanho Kim, Ashwin S. Shetty, Wen Yuan, Daniela Di Bella, Catherine Abbate, Paul Oyler-Castrillo, Xin Jin, Sean Simmons, Joshua Z. Levin, Juliana R. Brown, Paola Arlotta

**Author notes:** These authors contributed equally (Vahbiz Jokhi, Nuria Domínguez-Iturza, Kwanho Kim).

## Abstract

In the neocortex, oligodendrocytes produce distinct amounts of myelin in each cortical layer and along the axons of individual neuron types. Here we present a comprehensive single-cell molecular map of mouse cortical oligodendrocytes across different cortical layers and stages of myelination, spanning the initiation of cortical myelination into adulthood. We apply this dataset to show that neuron-class specific signals drive oligodendrocyte maturation and differential myelination across cortical layers. We find that each layer contains a similar compendium of oligodendrocyte classes, indicating that oligodendrocyte heterogeneity cannot explain layer-specific myelination. To evaluate whether neuronal diversity drives differential myelination across cortical layers, we generated a predicted ligand-receptor interactome between projection neuron types and oligodendrocyte states, across cortical layers and time. *In vivo* functional testing identified *Fgf18*, *Ncam1*, and *Rspo3* as novel, neuron-derived pro-myelinating signals. Our results highlight neuron-class-dependent control of myelin distribution in the neocortex.

## INTRODUCTION

Over the course of vertebrate evolution, the development of the myelin sheath has contributed to the expansion of the central nervous system (CNS) and the emergence of complex brain function. In the CNS, the lipid-rich myelin membrane is produced by oligodendrocytes (OL) that wrap the axons of neurons, forming myelinated segments (internodes), required to increase the velocity of action potentials and thereby enhance the efficiency of long-distance neuronal transmission^1,2^. Disrupted myelination can lead to many debilitating neurological disorders, including multiple sclerosis (MS) and schizophrenia^3,4^. Yet, our understanding of myelin formation and regulation at the molecular level is poor. The observation that oligodendrocytes are able to wrap myelin around paraformaldehyde-fixed axons and inorganic substrates^5–7^ has led to the idea that oligodendrocytes may be non-selective, and use physical cues such as axon caliber size as main determinants of myelination. However, the *in vivo* ensheathment of axons within neuronal networks requires precise patterning and spatial organization of myelin. *In vivo*, only axons become myelinated, not glial cells, vascular structures, neuronal dendrites, or cell bodies, and different neuronal subtypes show class-specific myelination patterns^8–11^.

In the cerebral cortex, the different layers present different amounts of myelination. This differential myelination is at least in part related to the distinct myelination density of the neuronal classes that occupy the different layers ^11,12^. We have previously shown that excitatory projection neurons (PNs) in different layers differ greatly in their longitudinal distribution of myelin^11^, with deep-layer neurons being generally more extensively and uniformly myelinated than upper-layer neurons. Similarly, Parvalbumin (PV^+^) interneurons of the cortex are more extensively myelinated than Somatostatin (SST^+^) or vasoactive intestinal polypeptide (VIP^+^) interneurons^8,9^, and display unique patterns of longitudinal distribution of myelin, contributing to layer variation in myelin content.

In theory, layer-specific differences in myelination could be driven by differential cues mediated by the different neuronal subclasses that populate each layer, or by intrinsic properties of OLs, such as OL heterogeneity^13,14^, developmental progression, or region-specific maturation^5,15^. Recent studies have identified cues that modulate the cell-intrinsic capacity of OLs to myelinate their targets^16^; for example, inhibitory signals that prevent myelination, such as Jam2, are expressed on neuronal somas and dendrites^7^. Moreover, mis-positioning of projection neuron subtypes from the deep layers to the upper layers results in increased ectopic myelination^11^, suggesting the presence of neuron-derived cues. Thus, the degree to which differential myelination is driven by OL heterogeneity vs layer-specific neuronal signals is unknown.

To understand the mechanisms of layer-specific myelination, we profiled OLs and their progenitors across cortical layers over a time-course of the early postnatal through adult mouse cortex, using single-cell RNA sequencing and spatial transcriptomics. We find that cortical oligodendrocytes differ primarily in maturation state rather than cell type within each layer. To test whether neuronal subtypes may differentially communicate with OLs to regulate their own myelination, we built an atlas of predicted ligand-receptor interactions between OL states and projection neuron subtypes. *In vivo* testing of candidate neuron-driven modulators of OPCs maturation and/or myelination showed that *Ncam1*, *Fgf18*, and *Rspo3* are capable of inducing ectopic myelination when expressed in upper-layer neurons. Our findings identify novel molecular signals, differentially expressed in specific neuronal populations, that mediate differential myelination across cortical layers.

## RESULTS

### Oligodendrocyte heterogeneity is due to differences in maturation state rather than oligodendrocyte subtypes in cortical layers

The cortex is organized into six radially-oriented layers which are marked by the presence of different projection neuron (PN) populations. Myelination in the cortex initiates in the deep layers around P10 and progresses into the upper layers over postnatal development^17–19^; however, the deep layers remain more highly myelinated at all ages^20,21^. This differential myelination could potentially reflect the presence of different OL populations across layers, or differential laminar distribution of mature myelinating OLs. Oligodendrocytes are heterogeneous, composed of transcriptionally distinct states^13,14^, but the nature and functional consequences of these states are unknown. To understand the mechanisms of layer-specific myelination, we profiled OLs and their progenitors across cortical layers over a time-course of the early postnatal through adult mouse cortex, using single-cell RNA sequencing (Figure 1A). We used a compound transgenic mouse line incorporating an inducible Cre driver and reporter (Fezf2*^CreERT^ x CAG^floxStop-tdTomato^*), which after tamoxifen injection at P3 resulted in the specific labeling of L5 corticofugal projection neurons (CFuPN) with tdTomato, crossed to the *Plp1-EGFP* reporter mouse, in which oligodendrocytes are labeled at all stages of development with EGFP. This genetic strategy allowed us to cleanly micro-dissect layers 1-4 (above the tdTomato^+^ cells), layer 5 (tdTomato^+^), and layer 6 (below the tdTomato^+^ cells) of the somatosensory cortex. We collected animals across a time course: i) early postnatal (postnatal day 7, P7), as OLs mature but before myelination begins, ii) juvenile (P14), when OLs begin to wrap axonal targets on L5 and L6 neurons, and iii) post-weaning and adult timepoints (P30 and P90), corresponding to myelination of deep-layer and upper-layer neurons, respectively (Figure 1A). Each time point included one male and one female mouse whenever possible (see STAR Methods). We dissociated and FACS-isolated Plp1-EGFP^+^ OL populations from each layer and age for scRNA-seq (Figure 1A and Figure S1A). In addition, in order to enrich specifically for the most mature populations of oligodendrocytes, we also isolated a separate sample at P90 containing only the cells with the highest EGFP intensity (“EGFP^+^-high”, Figure S1A; see STAR Methods). This approach allowed us to enrich for putatively mature oligodendrocytes, allowing a more detailed characterization of this population. Following the removal of neuronal and astrocytic cell populations and low-quality cells (Figure S1B-I) (see STAR Methods), the datasets together comprised a total of 38,496 cells (∼5000-14000 cells per time point; Table S1). Cells were clustered by a graph-based algorithm, and we assigned cell identities to the clusters based on the expression of known marker genes (Figure 1B-D and Figure S2A). We were able to identify populations corresponding to all developmental stages of OL maturation, namely progenitor states (OPC: OL precursor cells, COP: committed OL progenitors), premyelinating states (NFOL: newly-formed OL), and myelinating states (MFOL: myelin-forming OL, and MOL: Myelinating OL) (Figure 1B-D; see Tables S2 and S3 for a list of DE genes across oligodendrocyte types, ages and layers).

**Figure 1.**
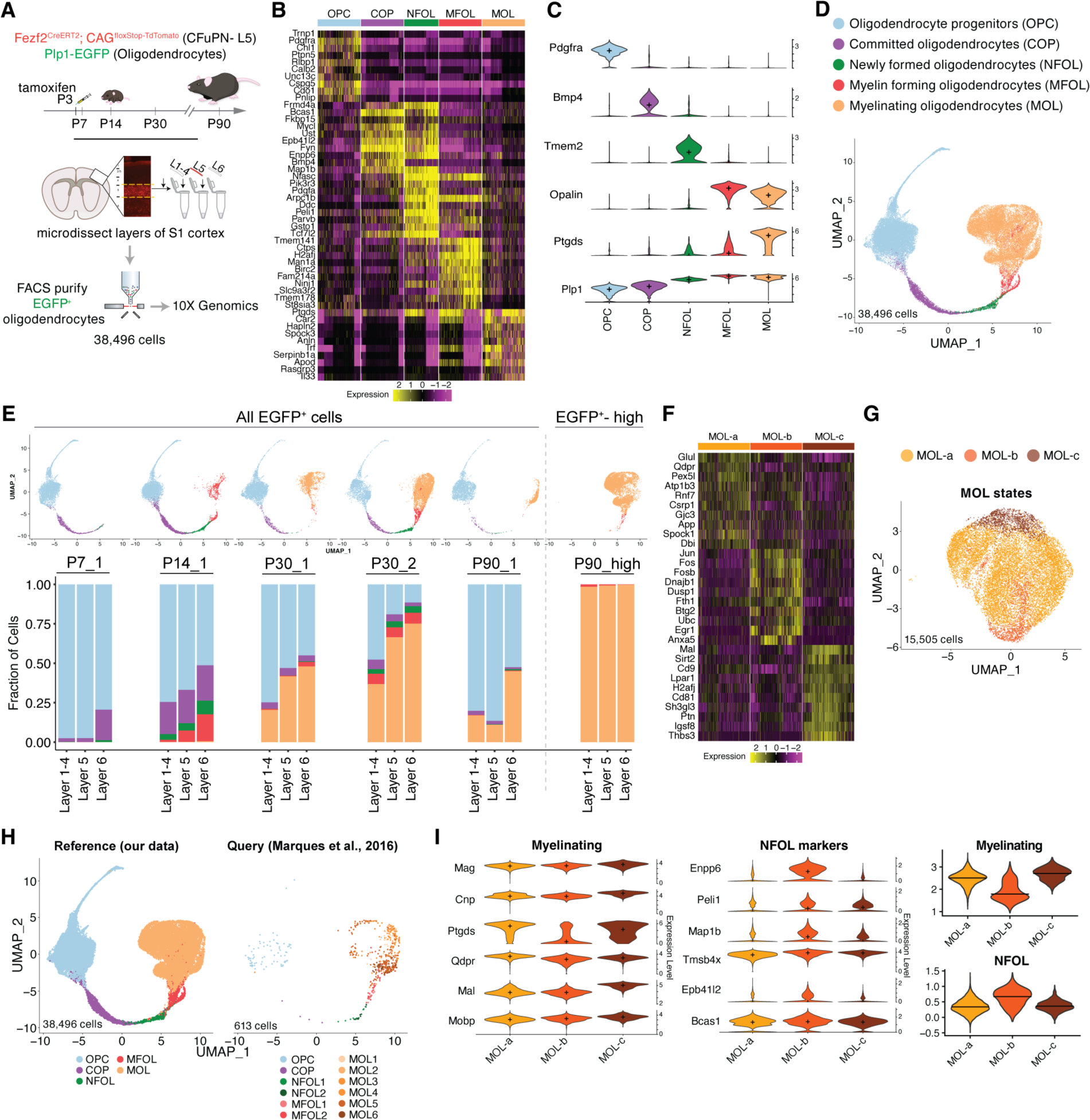
Single-cell profiling of cortical oligodendrocytes demonstrates positional heterogeneity of oligodendrocyte states. **(A)** Schematic of S1 cortex layer micro-dissection, EGFP^+^ OL isolation by FACS, and scRNA-seq from P7, P14, P30, and P90 *Plp1-EGFP* mice (pooled male and female for each age). **(B)** Heatmap of top 10 differentially-expressed genes for each OL population. **(C)** Violin plot of expression of key marker genes and *Plp1* across different OL populations. **(D)** UMAP of combined OL scRNA-seq datasets from micro-dissected layers, across all ages and experiments, color-coded by state. **(E)** Top: UMAPs of P7, P14, P30, P90 and P90_EGFP^+^-high scRNA-seq data, color-coded by OL state. Bottom: bar plot of P7, P14, P30, P90 and P90_EGFP^+^-high OL cluster proportions for each micro-dissected layer and experiment. Color-coding indicates the OL state. Data are normalized to the total cell count in each micro-dissected layer. **(F)** Heatmap of top 10 differentially-expressed genes for each MOL state. **(G)** UMAP of MOL subtypes, color-coded by the state identities identified in this study (MOL-a, MOL-b, and MOL-c). **(H)** UMAP of OLs from this study (left), and the projection of Marques et al. 2016 cortical OLs using Azimuth (right). **(I)** Left, violin plots of strongly-myelinating marker genes and NFOL marker genes in our MOL substates (color-coded). Right, violin plots of module score for myelinating and NFOL related genes in our MOL substates.

In the scRNA-seq dataset, we noted that, at all ages, the more myelinating OL populations were disproportionately present in the deep layers (Figure 1E). It is possible that differential myelination could be due in part to class diversity of mature myelinating OL populations. Prior studies have identified transcriptionally-distinct populations of MOLs^13^, but whether these represent different cell types or merely different cell states is unknown. To assess transcriptional diversity in mature OLs that may correlate to differential myelination across layers, we examined specifically the MOL population. We applied an unsupervised hierarchical clustering method to identify transcriptionally-distinct populations of MOLs in our data. This identified 3 subclusters, which we termed MOL-a, MOL-b, and MOL-c (Figure 1F and G). These populations were very similar to each other transcriptionally (Table S5); there were fewer than 100 DEGs (see STAR Methods), which differed mainly in the level of expression rather than binary absence or presence of expression. We then compared our dataset to previously-described cortical OL populations from Marques et al., 2016 ^13^; as that study sampled OLs from various brain regions, we used only the cortically-derived cells from their dataset, which were derived from the same region used here, the S1 somatosensory cortex (Table S1 and S5). We integrated our scRNA-seq results with the Marques et al. scRNA-seq dataset by projecting their cells onto our UMAP as a reference (Figure 1H). To compare our MOL clusters to the previously-described MOL populations, we used a random forest classifier to assign MOL identities from the Marques dataset^13^ to the MOLs in our dataset (see STAR Methods). We found that our MOL-a, MOL-b, and MOL-c showed only weak correlation to any of the Marques cortical mature MOL populations (Figure S2B-E), likely due to the different ages analyzed by the two studies and the comparatively low representation of cortically-derived MOLs in the Marques dataset (Table S5).

Assessing the expression of the genes related to OL maturation and myelination suggests that MOL-a and MOL-c represent more highly myelinating states, while MOL-b may represent a less mature myelinating MOL (Figure 1I). We examined the distribution of these three MOL clusters across ages and layers in our scRNA-seq dataset and found that the MOL-b population was more prevalent in L6 than in L1-5 (Figure S2F and G).

In order to more closely relate cell identities to their topographical organization in the cortex, we analyzed a previously-published *in situ* transcriptomics (Slide-seqv2) dataset of P56 somatosensory cortex^22^. For each of the projection neuron and oligodendrocyte cell types, we assayed the combined expression of the top differentially-expressed genes (DEGs) for that cell type (Figure 2A-D, Table S4). Expression of these gene sets in the Slide-seq dataset was found to be consistent with our scRNA-seq dataset: while OPCs and COPs are distributed across all cortical layers, maturing and mature OLs are predominantly present in the deep layers (Figure 1E, Figure 2E and F). Given that deep-layer neurons are more extensively myelinated at all ages, the data suggests that the spatial distribution of OL maturation states may be related to the local neuronal environment. We then examined the spatial distribution of each MOL cluster in the adult Slide-seq dataset, using a gene module score calculated from the top 10 DEGs per cluster, and found that all MOL states were more prevalent in the deeper layers: MOL-a, which is also the most abundant MOL subtype, was disproportionately present in L5, and MOL-b and MOL-c in L6 (Figure 2G-I). This suggests that the overall greater myelination observed in the deep layers may be associated with a higher density of mature myelinating OLs.

**Figure 2.**
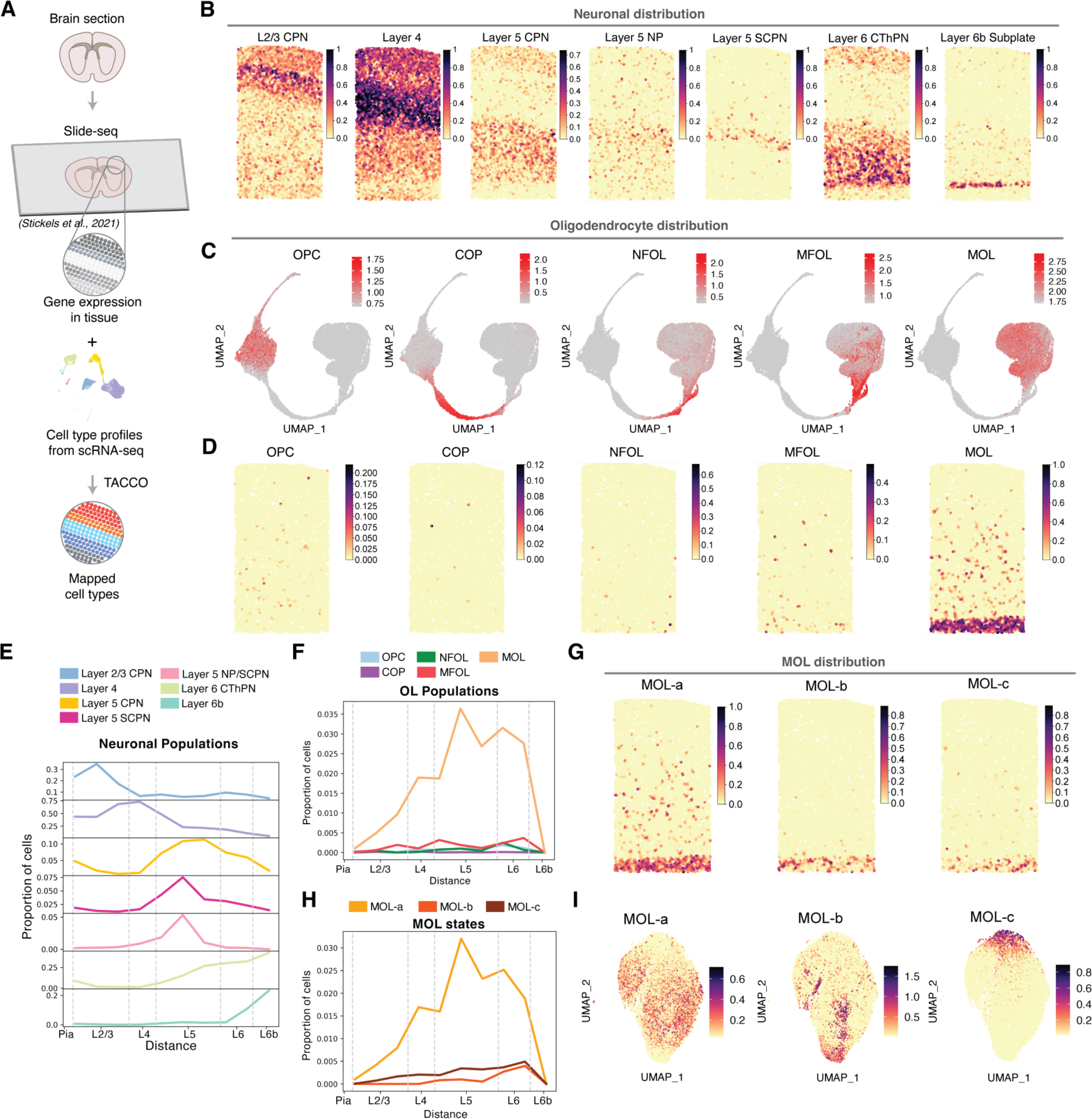
Spatial transcriptomics reveals positional heterogeneity of oligodendrocytes and MOL states. **(A)** Schematic summarizing the process of mapping the cell types found in our scRNA-seq study onto a matching spatial transcriptomic tissue previously published (Slide-seqV2)^22^ using TACCO^62^. **(B)** Spatial features plot of Slide-seqV2 of P56 somatosensory cortex, showing the module score of curated marker genes for each neuronal subtype (see Table S4), used to generate Figure 2E. **(C)** Gene module score of OL states (see Table S3) plotted onto a UMAP representation of the scRNA-seq data. **(D)** Gene module score of OL states plotted onto the spatial distribution of beads in the Slide-seq data, used to generate Figure 2F. **(E)** Spatial distribution of projection neuron types in a coronal section of P56 mouse cortex by Slide-seq. The graph shows the proportion of each cell type (y-axis) along various distances from the pia (x-axis). Dashed lines indicate cortical layers. **(F)** As in **E**, for OL populations (OPC, COP, NFOL, MFOL, and MOL). **(G)** Gene module score for the MOL states identified in this study (MOL a-c), plotted onto the spatial distribution of beads in the P56 somatosensory cortex Slide-seq data, used to generate Figure 2H. **(H)** As in **E** and **F**, spatial distribution of mature MOLs, classified by the state identities identified in this study (MOL-a, MOL-b, and MOL-c). **(I)** Gene module score for MOL states plotted onto the UMAP.

Our lab has previously shown that the density of myelination and the laminar location of mature oligodendrocytes correlates with the positioning of deep layer PN in young adult mice^11^. In order to understand whether myelin distribution across cortical layers was altered from the first stages of myelination, we examined whether mis-positioning of deep layer PN in upper cortical layers, at a juvenile age (P14) when upper layer PN are not yet myelinated, would influence myelination distribution in the cortex. We used the *Reeler* mouse model, which has a disorganized cortical lamination due to abnormal positioning of neuronal subtypes^23^ and which has been previously reported to have abnormal OPC proliferation and distribution, as well as myelination defects^24,25^. We performed immunohistochemistry in the somatosensory cortex of P14 mice; to quantify myelin distribution, we divided the cortex of wild-type (WT) and Reeler knock-out (*reln*^-/-^) mice from the pia to the white matter into horizontal bins, and quantified the fluorescence intensity of myelin basic protein (MBP), CTIP2 (layer 5/6 CFuPNs) and CUX1 (L2/3 upper layer CPNs) in each bin. In the control cortex, MBP intensity showed a strong correlation with CTIP2 signal (Pearson correlation, r = 0.963, P<0.0001), and a negative correlation with CUX1 signal (r = -0.67, P<0.01) (Figure S3, upper panels). In the *reln*^-/-^ cortex, despite disrupted laminar positioning of the neurons, we continued to observe a strong correlation of MBP with CTIP2 neurons (r = 0.77, P<0.05) and a negative correlation with CUX1 neurons (r = -0.74, P<0.01) (Figure S3, lower panels). The correlation of myelination density (MBP signal) with the local density of deep layer PNs (CTIP2^+^) suggests that the microenvironment created by these neuronal populations may regulate oligodendrocyte maturation and myelination.

### Class-specific neuron-oligodendrocyte molecular interactions

Given that the location of deep layer neurons can influence the distribution of myelin, we hypothesized that differential myelination is governed, at least in part, by a neuronal subtype-specific code of surface-bound or secreted molecules that affect oligodendrocyte maturation, myelin formation, and ultimately the distribution of myelin in the cortex. Neurons could in theory employ OL-facing signals such as inhibitory and/or attractive molecules to regulate differential myelination (Figure 3A). Thus, an understanding of molecules differentially expressed in weakly-vs. highly-myelinated PN subtypes could enable the identification of signals that play a role in neuron-OL communication.

**Figure 3.**
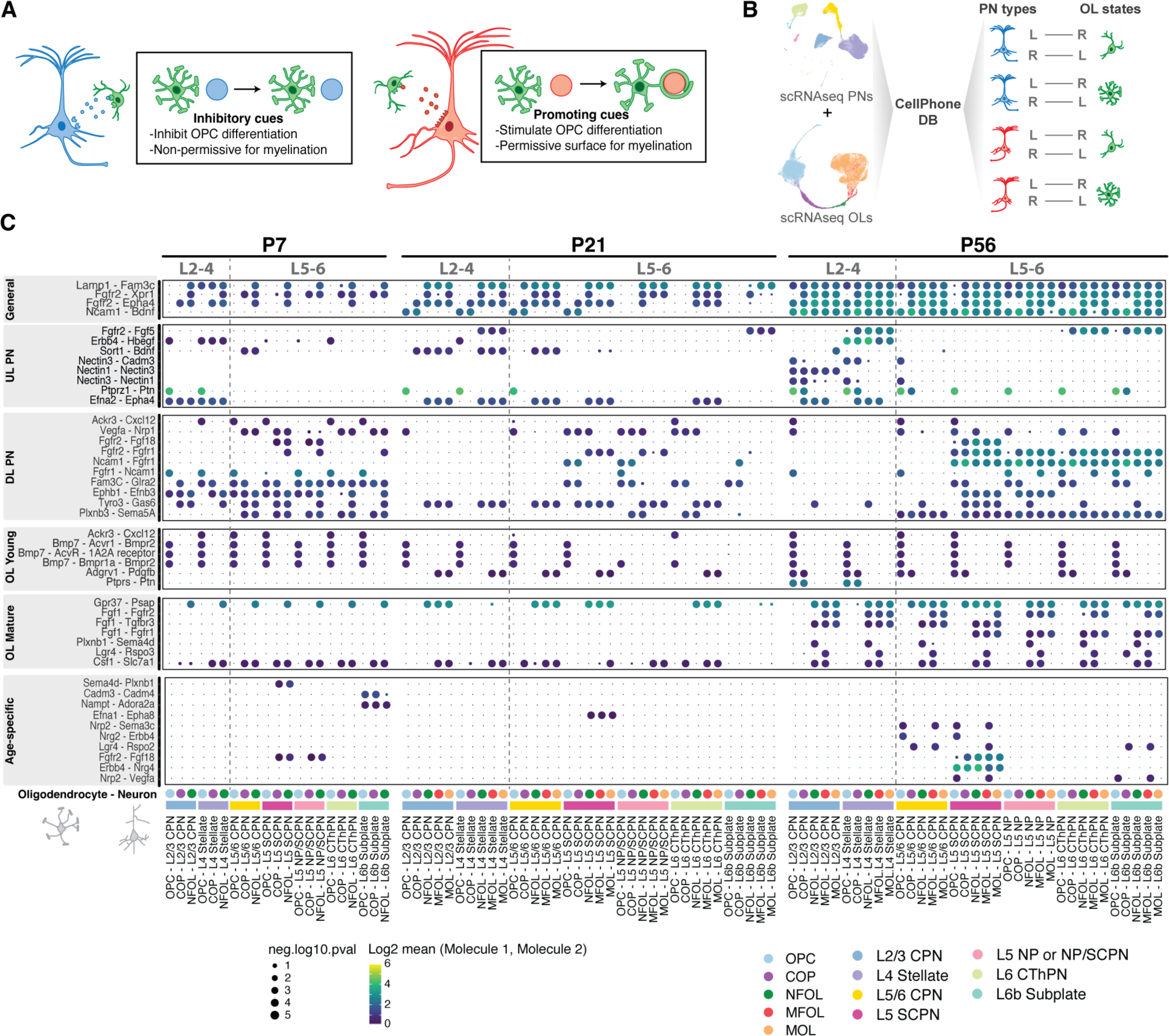
Interactome of pyramidal neuron-oligodendrocyte communication in the neocortex. **(A)** Schematic of hypothesized Ligand-Receptor (L-R) interaction. Neurons producing inhibitory (blue) or attractive (red) cues and oligodendrocytes (green) are represented. **(B)** Schematic summarizing the generation of the predicted interactome using transcriptomic data from projection neurons (PNs) and oligodendrocytes (OLs) as input to CellPhone DB to identify predicted ligand (L) and receptor (R) pairs. **(C)** Dot plot of L-R expression between PN subtypes and OL states at P7 (left), P21 (middle), and P56 (right). L-R pairs that pass significance threshold are shown for each selected grouping, based on cell-type specificity and age. On the x-axis, PN subtypes are represented in colored rectangles, and OL states are shown in colored circles. The size of each data point represents -log_10_ adj. *P* values, and the color indicates the average log_2_ mean expression level of the interacting molecules from the two participating cell populations. L-R pairs that are not significant are represented with the smallest circle, while those that are not expressed have no circle. The label for each row gives the molecule expressed in the oligodendrocyte population followed by the molecule expressed in the neuronal population.

Given that developmental myelination is a protracted process and continues well into adulthood, we examined neuron-OL interaction over an extended time course. We generated single-cell sequencing datasets from WT P7 and P21 whole somatosensory cortex, and identified all cell types present (Figure S4). In addition, we also analyzed a publicly-available scRNA-seq dataset from the primary somatosensory (S1) cortex at P56 from the Allen Brain database^26^. Within these datasets, we extracted all PN profiles and oligodendrocyte states, and classified them based on gene expression and laminar location (Figure S5).

We explored Ligand-Receptor (L-R) interactions between PN subtypes and oligodendrocyte states by employing a curated molecular database, CellPhoneDB^27^, with stringent cutoffs (see STAR Methods). To identify proteins expressed across different PN subtypes that have cognate ligands or receptors on the different OL states, PN subtypes were tested against the different states of oligodendrocytes in the same scRNA-seq datasets for each of the 3 ages. Analysis of these datasets resulted in >120 L-R pairs with significant expression in at least one OL-PN pairwise test (Benjamini Hochberg adj. *P* < 0.05; Table S6, Figure 3B and C).

The results included well-established signaling molecules that control oligodendrocyte differentiation, such as *Gas6-Tyro3* and *Cxcl12-Ackr3*. *Gas6* stimulates oligodendrogenesis and myelination, and loss of *Gas6* is associated with loss of oligodendrocytes^28^. *Cxcl12* promotes OPC differentiation *in vitro*^29^ and remyelination *in vivo* in the adult CNS^30^. The presence of these expected interacting candidate molecules adds validity to the other predicted interactions, a large number of which have not been previously implicated in myelination.

We identified candidate L-R interactions predicted to involve specific cell populations and ages: i) interactions of most or all PN subtypes with all oligodendrocytes (termed “general”), ii) interactions with upper layer PNs (UL PN), iii) interactions with deep layer PNs (DL PN), interactions specific to either iv) young or v) mature oligodendrocytes, and vi) interactions specific to a particular age. To narrow down possible targets, we focused on those interactions where the candidate molecules were differentially expressed between upper- and deep-layer neurons, as these are the most likely to represent candidates that could mediate differential distribution of myelination across layers. We found several genes that were differentially expressed between the UL and DL neurons, either at a specific age or independently of age (Figure 3C, Figure 4A and B, Figure S6). Interestingly, *Gas6* was differentially expressed across PNs, being abundantly present in deep layer PNs at all ages, while Cxcl12 was predominantly expressed in DL PNs only at younger ages (P7 and P21) (Figure 4A and B). Other genes, previously unclassified as myelinogenic, such as *Fgf18*, *Ncam1*, *Sema5a* or *Glra2*, had a higher expression in DL PNs at all ages, and therefore represent possible candidates to promote layer specific myelination. Conversely, genes such as *Ptn* or *Epha4* had greater expression in UL PNs, and thus may represent candidate molecules that prevent myelination in upper cortical layers. Interestingly, genes such as *Nectin1/3* or *Xpr1* had layer-specific expression only at certain postnatal stages, while *Bdnf* changed the expression pattern across time (Figure 4).

**Figure 4.**
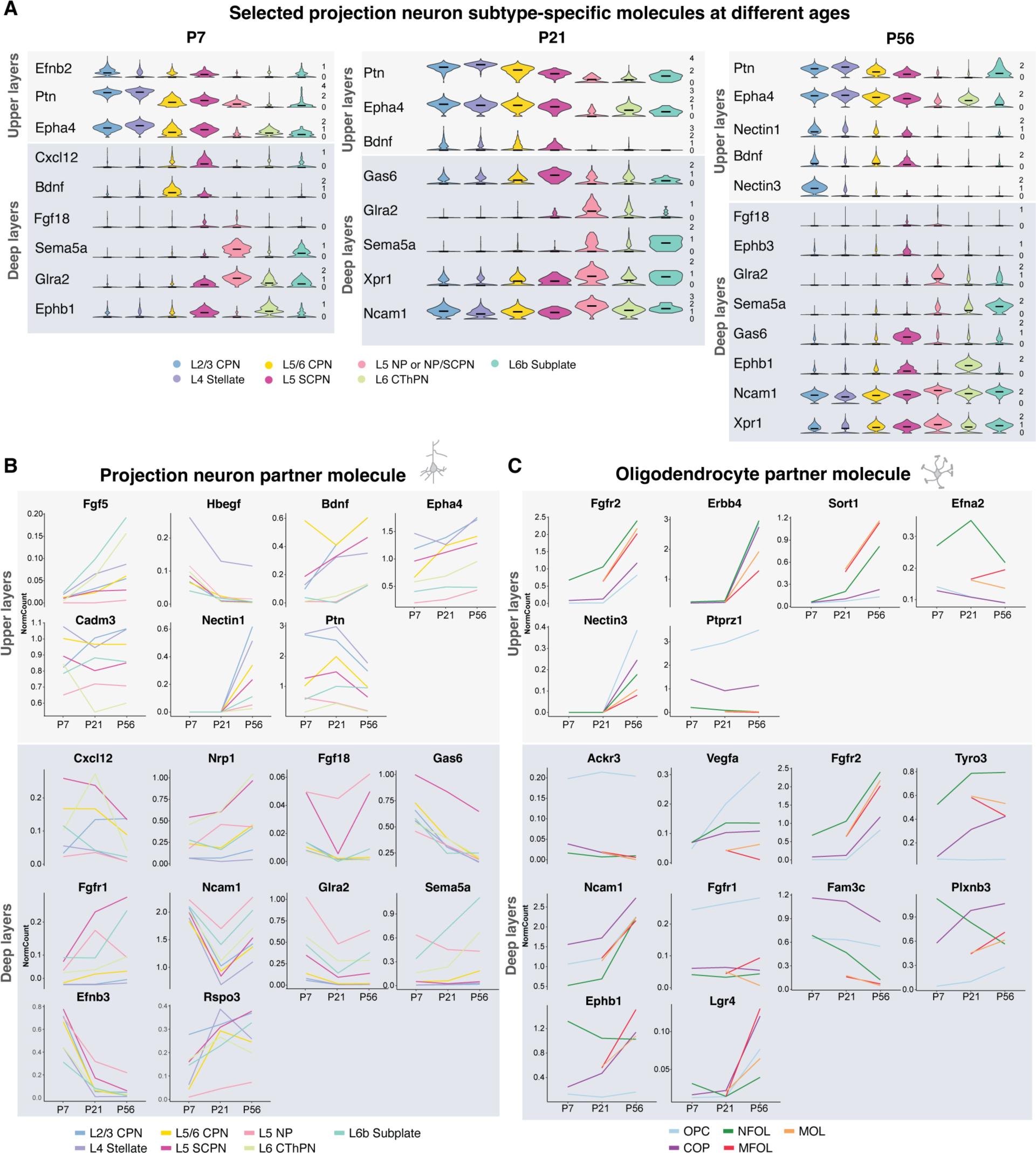
Ligand and receptor pairs show specificity across layers and ages. **(A)** Violin plots of normalized gene expression level (in natural log scale) among PN subtypes from scRNA-seq data at P7 (left), P21 (middle), and P56 (right). PN subtypes are color-coded. **(B)** Average expression levels over time of selected candidate genes from L-R interaction analysis in neurons. Neuron types are color-coded. Up, genes with higher expression in UL-PN. Down, genes with higher expression in DL-PN and/or more mature substates of oligodendrocytes. **(C)** Expression levels in oligodendrocytes of the cognate ligand or receptor for the molecules in B.

Altogether, these genes represent candidate effectors of a subtype-specific molecular code that mediates differential OL maturation and / or myelination in the upper vs deep cortical layers.

### *In vivo* screen of novel molecular mediators of layer-specific myelination in the neocortex

To test the functional effects of our candidate L-R genes on differential myelination, we applied an *in vivo* overexpression screen to identify candidates that promote myelination. We selected candidates that were more highly expressed in deep-layer neurons (CFuPNs), and which may thus represent “inductive” signals, and overexpressed them in upper-layer neurons (L2/3 CPNs) via *in utero* electroporation at E14.5. We used immunofluorescence to examine increased OL maturation (APC, CC1 clone) and myelination (myelin basic protein, MBP) near the site of electroporated, overexpressing neurons in juvenile animals (P21) (Figure 5A). Given that myelination in the cortex varies with rostro-caudal and medio-lateral position, we used the corresponding region of the un-electroporated contralateral hemisphere as control. We note that a limitation of this approach is that contralateral axonal projections of the electroporated L2/3 CPNs, which also overexpress the target gene, could influence myelination in the contralateral region, and thereby preclude the detection of subtle changes.

**Figure 5.**
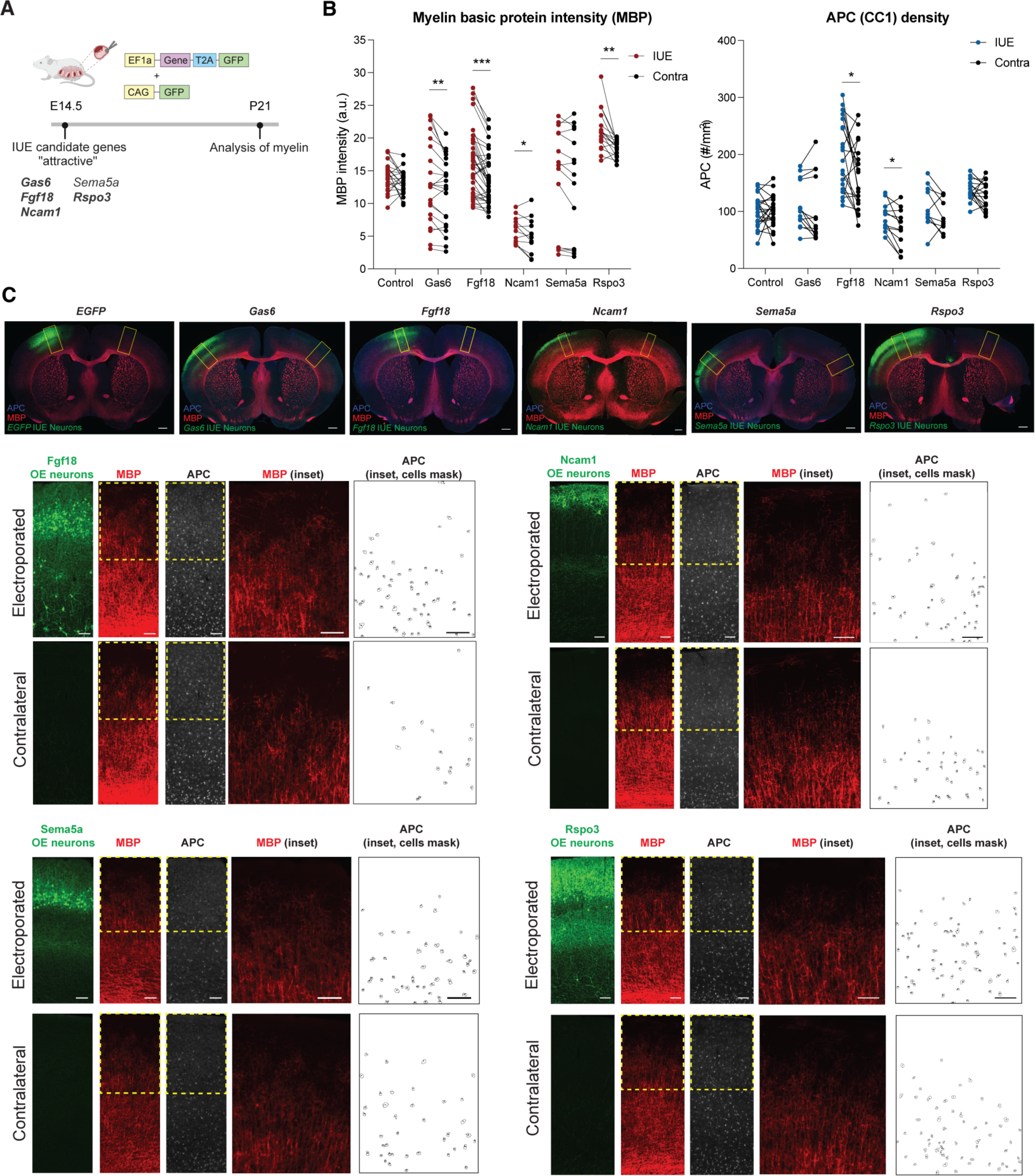
*In vivo* screen of candidate mediators of myelination. **(A)** Schematic of experimental setup. In-utero electroporation (IUE) was performed at embryonic stage 14.5 (E14.5) with an expression vector containing the target gene of interest and the pCAG-EGFP vector. Progeny was analyzed by immunohistochemistry at postnatal day 21 (P21). Genes tested are listed, with significant hits in bold. **(B)** Quantification of myelin basic protein (MBP) intensity (left) and APC^+^ (CC1, mature OL marker) cell density (right) of electroporated (IUE, red dots for MBP and blue dots for APC) and contralateral (contra, black) cortical regions at P21 after in utero electroporation (IUE) of candidate mediators of myelination at E14.5. Each data point represents one brain section (MBP: *EGFP* n=5 mice, N=23 sections; *Gas6* n=3 mice, N=22 sections; *Fgf18* n=4 mice, N=39 sections; *Ncam1* n=3 mice, N=12 sections; *Sema5a* n=5 mice, N=15 sections, *Rspo3* n=5 mice, N=34 sections. APC: *EGFP* n=5 mice, N=23 sections; *Gas6* n=3 mice, N=13 sections; *Fgf18* n=4 mice, N=20 sections; *Ncam1* n=3 mice, N=12 sections; *Sema5a* n=3 mice, N=10 sections, *Rspo3* n=5 mice, N=34 sections) (p-values from linear models, MBP: *EGFP* p=0.0589, *Gas6* p=0.006, *Fgf18* p=0.000009, *Ncam1* p=0.048, *Sema5a* p=0.053, *Rspo3* p=0.0023; APC: *EGFP* p=0.986, *Fgf18* p=0.043, *Ncam1*p=0.025, *Sema5a* p=0.213, *Rspo3* p=0.218) (* p<0.05; ** p<0.01; *** p<0.001). Lines connecting two data points represent corresponding IUE and contralateral regions of each section. **(C)** Top, representative section of IUE cortex for control (*EGFP*) and overexpressed candidates (*Gas6*, *Fgf18*, *Ncam1*, *Sema5a* and *Rspo3*). Sections were immunolabelled for MBP in red and APC (CC1, mature oligodendrocyte marker) in blue. Electroporated cells express EGFP. Analyzed electroporated and contralateral regions are highlighted with a yellow box (scale bar: 500µm). Bottom, representative images of IUE and contralateral P21 S1 cortical regions analyzed for MBP intensity (red) and APC density (blue). Over-expressed candidate genes (*Fgf18*, *Ncam1*, *Sema3a*, and *Rspo3*) are indicated for each panel. Inset: magnification of upper cortical layers, showing MBP intensity and APC count (scale bar: 100µm).

We first tested a known modulator of myelination, *Gas6*, as a positive control^28^. As expected, *Gas6* overexpression led to increased myelination in the electroporated region (P<0.01) (Figure 5B and C, Figure S7A and B). The empty EGFP expression vector (negative control) resulted in no difference in myelination or oligodendrocyte maturation between electroporated and contralateral cortical regions (Figure 5B and C, Figure S7A and B). We tested four predicted “inductive” candidates from the interactome; three of the candidates (*Fgf18*, *Ncam1,* and *Sema5a*) are enriched in deep layer neurons (Figure 3C, Figure 4), and while the expression of the fourth candidate, *Rspo3,* is similar in all PN sub-types, its ligand (*Lgr4*) is specifically enriched in mature oligodendrocytes (Figure 3C, Figure 4). Three of the candidates, *Fgf18*, *Ncam1,* and *Rspo3,* significantly increased myelination in the electroporated region compared to the contralateral hemisphere (linear model, *Ncam1* p<0.05, *Fgf18* p<0.001, *Rspo3* p<0.01; see STAR Methods); the fourth molecule, *Sema5a*, approached but did not reach significance (p=0.053). Two candidates, *Fgf18* and *Ncam1*, were also able to promote OPC maturation (linear model P< 0.05 *Fgf18*, P<0.05 *Ncam1*) (Figure 5B and C, Figure S7B). FGF18 has been shown to be a mitogen that promotes proliferation of OPCs^31^, and *in vivo*, activation of one of its receptors, FGFR2, regulates myelin growth^32,33^. Importantly, *Ncam1* and *Rspo3* have not been previously implicated in myelinogenesis, and are novel candidates for neuron class-driven myelination in the neocortex. We next calculated a gene module score for the three receptor genes expressed by oligodendrocytes (*Lgr4*, *Fgfr1,* and *Fgfr2*) that correspond to these three candidate ligands: in our scRNA-seq dataset, we found that the module score increased with age, and was most highly expressed in oligodendrocytes of the deep layers (Figure S7C).

Given the small effect size of our candidate genes in stimulating OPC maturation and myelination, and the fact that several candidates are able to modulate myelination, we hypothesized that differential myelination is likely mediated by more complex gene regulatory networks. Therefore, we used co-electroporation to co-express the two candidate genes which promoted both myelination and OPC maturation, *Fgf18* and *Ncam1*. Indeed, co-electroporation at E14.5 and subsequent examination at P14 (Figure 6A, Figure S7D) and P21 (Figure 6B, Figure S7D) demonstrated a strong synergistic effect of the two genes for both increasing OPC maturation (as denoted by increased APC^+^ cell density) and myelination (increased MBP intensity). Interestingly, StringDB, a curated database of all protein-protein interactions^34^, indicates that both the FGFR1 and FGFR2 receptors can respond to either of these two ligands. This further support a model by which the activation of the FGF pathway stimulates oligodendrocyte maturation and myelination in the vicinity of deep layer neurons, which express these ligands more abundantly (Figure 3 and 4).

**Figure 6.**
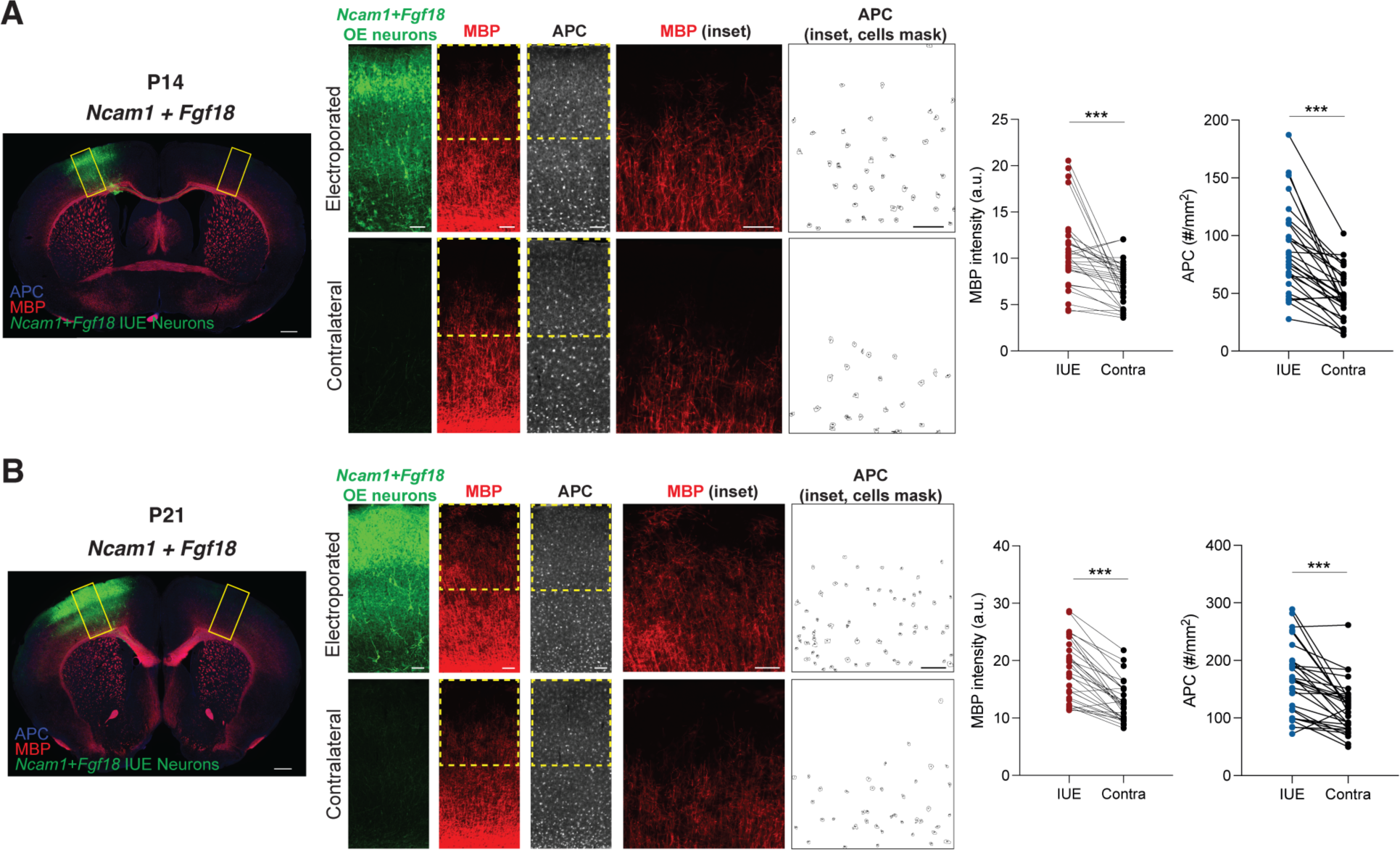
Synergistic effect of *Fgf18* and *Ncam1* overexpression on myelination. **(A)** Left, representative section of IUE cortex at P14 overexpressing *Fgf18* and *Ncam1* together. Sections were immunolabelled for MBP (red) and APC (blue). Electroporated cells express EGFP. Analyzed electroporated and contralateral regions are highlighted with a yellow box (scale bar: 500µm). Center, representative electroporated and contralateral S1 cortical region at P14 after IUE at E14.5 of combined *Fgf18* and *Ncam1* (scale bar: 100µm). Inset: magnification of upper cortical layers, showing MBP intensity and APC count (scale bar: 100µm). Right, quantification at P14 of myelin basic protein (MBP) intensity and APC^+^ cell density in upper layers of electroporated (IUE, red for MBP and blue for APC) and contralateral (contra, black) cortical regions after IUE. Each data point represents one brain section (n=5 mice, N=30 sections total). (p-values from linear models: Total: MBP p=0.0006; APC p=0.0000008) (* p<0.05; ** p<0.01; *** p<0.001). **(B)** As in **A**, but at P21. Each data point represents one brain section (n=5 mice, N=30 sections total) (p-values from linear models, Total: MBP p=0.00000001; APC p=0.00004) (* p<0.05; ** p<0.01; *** p<0.001).

Taken together, our results indicate that different cortical projection neuron classes use specific molecular codes to communicate with oligodendrocytes, ultimately guiding differential myelination across cortical layers. Our screen allowed us to identify *Fgf18* and *Ncam1* as novel myelination-stimulating candidate genes produced preferentially by PNs of the deep layers, pointing at new pathways underlying differential OL maturation and myelination in the different cortical layers and the role of PN diversity in developing myelin maps in the neocortex.

## DISCUSSION

Different regions of the CNS are known to be differentially myelinated, but the mechanisms that control that diversity are poorly defined. To investigate cortical myelination diversity, here we generated a single-cell molecular map of cortical oligodendrocytes from different layers and stages of myelination, and created a predicted ligand-receptor interactome between projection neuron classes and oligodendrocyte states across postnatal development. We show that the composition of PN subtypes in the different cortical layers is a determinant of myelin density and OL state distribution. This adds to the evidence that projection neuron diversity in the neocortex forms the framework for instructing the localization, states, and function of other cell types, including interneurons and microglia^35–37^, and now oligodendrocytes (this study). These findings put forward the principle that neuronal diversity in the CNS is not only important *per se*, but also drives interactions with glia to develop higher-order features of the brain, including layer-specific myelination.

The hypothesis that projection neuron subtypes use differential signaling to influence oligodendrocyte-lineage cells around them led us to profile different projection neuron populations to define the signals by which they act. Functional testing of these candidate genes allowed us to identify three novel pro-myelinating proteins, NCAM1, FGF18, and R-spondin-3, which could guide differential myelination in the cortex (Figure 5). Of relevance for therapeutic research, co-expression of NCAM1 and FGF18 resulted in a strong synergistic effect on OL maturation and myelination, suggesting that myelination may be better controlled by a combinatorial code of signals.

A clinical goal in demyelinating disorders has been to promote efficient myelin repair by directly promoting OL differentiation and remyelination^38^. Some demyelination phenotypes appear to be associated with a lack of myelination activity rather than a lack of OLs^39,40^: for example, multiple sclerosis lesions in post-mortem brains still contain mature OLs, yet there is a failure to remyelinate axons. Interestingly, the unsialylated form of NCAM1, one of the molecules we found to promote myelination, is enriched in the cerebrospinal fluid of multiple sclerosis patients that respond well to steroid therapy compared to those that do not^41,42^, suggesting that increased NCAM1 levels could be promoting remyelination in those patients. Over normal development, NCAM1 in the brain transitions from a highly-polysialylated embryonic form to a less-polysialylated adult form^43^ which changes its binding properties^44^. Interestingly, polysialylated NCAM1 (PSA-NCAM) reduces OPC maturation and prevents myelination^45^, and the onset of myelination in the human fetal forebrain coincides with the down-regulation of PSA-NCAM^46^. Together, this suggests that regulation of NCAM1 sialylation may be an important factor in the control of myelination.

Research over recent years demonstrated that many neuropsychiatric conditions are characterized by white matter abnormalities^4,47^. Interestingly, defects in many of the novel myelinogenic molecules we identified in this study have been implicated in neuropsychiatric conditions. SNPs in *NCAM1* contribute to differential risk of developing bipolar disorder and schizophrenia^48^, and, recently, autoantibodies against NCAM1 were found in patients with schizophrenia^49^. Moreover, disruption in the Wnt and FGF signaling pathways, which are activated by *Rspo3* and *NCAM1* / *Fgf18*, respectively, have been observed in patients with schizophrenia, bipolar disorder, and major depressive disorder^50,51^. While defects in NCAM1 or the FGF signaling pathway have not been reported to be directly associated with myelination abnormalities in patients, our findings raise the hypothesis that defects in those pathways may lead to impaired myelination, contributing to the clinical phenotypes.

All together, our results indicate that different cortical projection neuron classes use specific molecular codes to communicate with oligodendrocytes, ultimately guiding differential distribution of myelin in the cortical layers. A more comprehensive understanding for how different classes of neurons in the brain affect OL biology and myelination, will in the future increase our understanding of both brain development and of myelin dysfunction in disease states.

## STAR METHODS

### Resource availability

### Lead contact

Further information and requests for resources and reagents should be directed to and will be fulfilled by the lead contact, Paola Arlotta.

### Materials availability

This study did not generate any new unique reagents.

### Experimental model and subject details

### Mice and husbandry

For FACS-isolation of oligodendrocyte-lineage cells from different cortical layers, we used the following mouse lines: *Plp1-EGFP*^52^, *Fezf2^CreERt2^* ^53^, conditional reporter line *CAG^floxStop-tdTomato^ Ai14-Gt(ROSA)26Sor^tm9(CAG-tdTomato)Hze^*(IMSR Cat# JAX:007909, RRID:IMSR_JAX:007909)^54^. Mice from these different lines were crossed to generate *Fezf2^CreERT2/+^; td-Tomato^+/-^; Plp1-EGFP^+/-^* mice to simultaneously label oligodendrocytes and corticofugal projection neurons (CFuPN). Mice were injected at postnatal day 3 with 4-hydroxytamoxifen (Sigma H6278) in corn oil (single dose of 100 mg tamoxifen per kg body weight). For PN mis-localization experiments, *Reeler* KO (*Reln^-/-^* RRID:IMSR_JAX:000235) mice were used. Transgenic mice were maintained on a C57BL/6J background, and both male and female mice were used for experiments. *In utero* electroporation procedures were performed on CD-1 timed pregnant females purchased from Charles River Laboratories. All procedures were designed to minimize animal suffering and approved by the Harvard University Institutional Animal Care and Use Committee (IACUC) and performed in accordance with institutional and federal guidelines. Mice were housed in individually ventilated cages using a 12-hour light/dark cycle and had access to water and food *ad libitum*.

### Method details

### Cortical microdissection, single cell suspension and FACS

#### Microdissection of layers from S1 cortex (for scRNA-seq for OL)

For each sample, one male and one female *Fezf2^CreERT2/+^; td-Tomato^+/-^; Plp1-EGFP^+/-^* mouse at each time point were used for S1 cortex layer microdissections. Mice at different time points (P7, P14, P30, and P90) were anesthetized with isoflurane, followed by quick decapitation, and brains were dissected and transferred into ice-cold low-fluorescence Hibernate-A (Brain Bits, HALF 500). Brains were coronally sectioned at 200-400 µm in Hibernate-A on a Lecia vibratome, and sections were selected exclusively from the S1 cortex: specifically, from the point where the corpus callosum crosses the midline (bregma +1.1mm) to the point where the hippocampus dips below the dentate gyrus (bregma -1.94mm). Using a fluorescence dissecting microscope, the S1 region was further micro-dissected in cold Hibernate-A into 3 layer compartments (layers 1-4, L5 and L6) using Fezf2-tdTomato^+^ cells to demarcate L5. Care was taken to exclude the corpus callosum since it is known to have differential OLs from the cortex^55^.

#### Cortical dissection (for scRNA-seq of all cell types)

Somatosensory and motor cortex from wild-type animals was dissected at P7 and P21. Mice were anesthetized with isoflurane, followed by quick decapitation, and brains were dissected and transferred into ice-cold media. The somatosensory and motor cortex were excised and transferred to fresh ice-cold media.

#### Preparing single-cell suspensions

Live single-cell suspensions were prepared for fluorescence activated cell sorting (FACS) using an adapted protocol from Worthington Papain Dissociation System kit (Worthington Biochemical, LK003150)^56^. Briefly, micro-dissected brain tissues were finely chopped using a blade and incubated in EBSS with Papain and DNAse for 30 mins (P7) to 1.5 hours (P90) at 37°C. This was followed by gentle trituration to break up the tissue into a single-cell suspension, and papain activity was blocked using the ovomucoid inhibitor. The dissociated cells were resuspended in ice-cold Hibernate-A containing 10% fetal bovine serum (FBS, SH30070.03HI).

### Fluorescence-activated Cell Sorting (FACS)

#### Oligodendrocyte purification

Plp1-EGFP^+^ oligodendrocytes were sorted on a MoFlo Astrios EQ Cell Sorter (Beckman Coulter) using a 100 μm nozzle. For most samples, all EGFP^+^ populations of different EGFP intensity were collected, since they likely represent different stages of OL maturation. For P90, we collected both this all-EGFP^+^-cells sample, and a second sample (from a different biological replicate) gated on the population with the highest GFP signal (labelled “P90_EGFP^+^-high”), with the aim of enriching for mature oligodendrocytes to allow detailed characterization. On average, 5,000 to 8,000 EGFP^+^ cells were sorted per layer and age. The sorted cells were spun down at 200g G for 5 min and resuspended in PBS containing 0.04% BSA (NEB B9000S).

#### All cortical populations

Live cells were isolated by sorting on a MoFlo Astrios using a 100μm nozzle, as DAPI-negative and Vybrant DyeCycle Ruby (Thermo Fisher V10273)-positive events.

### Single-cell sequencing

#### Oligodendrocytes at P7, P14, P30 and P90

Approximately 6-8,000 sorted EGFP^+^ oligodendrocytes were loaded onto a Chromium™ Single Cell 3’ Chip (10x Genomics, PN-120236) and processed through the Chromium Controller to generate single-cell gel beads in emulsion. Single-cell RNA-Seq libraries were prepared with the Chromium™ Single Cell 3’ Library & Gel Bead Kit v2 (10x Genomics, PN-120237) or v3 for the second experiment at P30 and P90 (10x Genomics, PN-1000121). Libraries from different layers were pooled based on molar concentrations and sequenced on a NextSeq 500 instrument (Illumina) to an average read depth of 30,000 reads per cell.

#### All cell populations at P7 and P21

Approximately 10,000 cells from dissociated cortexes were loaded onto a Chromium™ Single Cell 3’ Chip (10x Genomics, PN-120236) and processed through the Chromium Controller to generate single-cell gel beads in emulsion. Single-cell RNA-Seq libraries were prepared with the Chromium™ Single Cell 3’ Library & Gel Bead Kit v2 (10x Genomics, PN-120237). Libraries from different replicates were pooled based on molar concentrations and sequenced on a NextSeq 500 instrument (Illumina) to an average read depth of 30,000 reads per cell.

The read distribution was as follows: 26 bases for read 1, 57 bases for read 2 and 8 bases for Index 1. A sequencing saturation of 70-80% was targeted for all samples.

### scRNA-seq Data Analysis Pipeline

Single-cell RNA-seq binary cell call (BCL) files for experiments from P7, P14, P30 and P90 oligodendrocytes and all cell types from P7 and P21 mice were converted to FASTQs using the mkfastq function from Cell Ranger software version 3.0.1 for oligodendrocytes and 3.0.2 for all cell types, with the default parameters. FASTQs were aligned to the mm10 (Ensembl 93) reference transcriptome, and gene expression matrices were obtained using the Cell Ranger count function of with the default parameters. CellBender^57^ was then applied to remove ambient RNA (remove-background method with epochs=150) and Scublet^58^ to compute doublet score for each cell (expected_doublet_rate=0.05, min_counts=2). R version 4.0.3 and Seurat package version 4.0.0^59^ were used to perform all downstream analyses.

For initial quality filtering, genes expressed in more than three cells were kept, and cells expressing more than 500 genes were kept. Cells with greater than 10% mitochondrial gene expression (percent.mt) were removed. The SCTransform function from Seurat (sctransform package version 0.2.1) with vars.to.regress parameter (to account for batch effects) set to percent.mt and sample (two biological replicates for each micro-dissected cortical layer) was used to normalize, find variable genes, and scale the dataset. The identified list of variable genes was used to perform principal component analysis (RunPCA function with the default parameters). Cell clusters were identified using the top 30 principal components and Louvain clustering algorithm (FindNeighbors function with dims=1:30 and FindClusters function with resolution=0.5 unless otherwise indicated). For visualization, the top 30 PC dimensions were further reduced to two uniform manifold approximation and projection (UMAP) dimensions (RunUMAP function with dims=1:30). The procedure, from SCTransform to dimensionality reduction for visualization, was repeated each time unwanted cell populations were removed, allowing more-accurate capture of the variation among the remaining cells, until we were left with only the cells of interest. Visualization of scRNA-seq was performed with ggplot2 (version 3.2.1, https://ggplot2.tidyverse.org)^60^.

Cluster-specific differentially expressed genes (DEGs) were identified by fitting a negative binomial generalized linear model (glm.nb function from MASS R package v7.3-53.1) to each gene while considering the differences in total number of UMIs per cell. The p-value for each gene was obtained by Wald test. The resulting list of DEGs were filtered to have adjusted p-value less than 0.05.

#### Oligodendrocytes Across Ages

The two experiments went through separate QC, because they were processed at different times using different 10X chemistry kit versions. For replicate1, the initial cell clustering identified 17 different clusters. Clusters with high expression of neuron-, microglia- or astrocyte-specific genes were removed (211 excitatory neurons with *Neurod2, Satb2,* and *Cux1*; 30 inhibitory neurons with *Gad1* and *Gad2*; 31 microglia with *Aif1*; 1313 Astrocytes with *Aqp4, Aldh1l1*). Clusters with low-quality signals (low UMI/cell, low genes/cell and high percent.mt; 144 cells) were also removed. For the second experiment, we only found one cluster of Astrocytes (109 cells), which was removed.

After QC, the remaining OLs (22189 from experiment 1 and 16307 from experiment 2) were merged into the same Seurat object and processed together. We performed Reference Component Analysis-based analysis (code available on Github), using a previously-published dataset ^13^ as a reference to co-embed oligo substates (OPC, COP, NFOL, MFOL and MOL) from different ages. In this reduced dimension, new clusters were identified and annotated with OL type by the DEGs.

The MOL population was isolated and processed to further identify subtypes. A new set of variable genes was found, and sample-specific covariates (animal and cortical layer) were regressed out when scaling the data. For batch correction, Harmony (R package v1.0) was applied to age. To evaluate the significance of MOL sub-clustering, we applied the testClusters function from scSHC (R package v0.1.0)^61^, reducing the initial seven clusters (FindCluster function from Seurat with resolution=0.3) to three significant ones. We noted that some of the genes with highest effect size among sub-clusters were immediate early genes, which could be contributing to the clustering.

#### All cell types across ages

Initial cell clustering identified 32 and 34 different clusters in P7 and P21 datasets, respectively. Clusters with high doublet scores and those low in quality (low UMI/cell, low gene/cell, and high percent.mt) were removed (2803 cells in P7, and 1272 cells in P21). After processing the remaining cells for each age (36,898 cells for P7, and 53,732 cells for P21), the identified cell clusters were assigned cell type labels based on their cluster-specific marker genes (Supplemental Table 8). Clusters that expressed markers of glutamatergic neurons (ExN; *Slc17a7* and *Neurod2)* and oligodendrocytes (OL) were further processed to annotate the subtypes. ExN and OL were individually processed to yield cell clustering. Clusters were then assigned a subtype label (ExN subtypes: UL_CPN, L4_Stellate, DL_CPN, NP, SCPN, CThPN, and Subplate; OL subtypes: OPC, COP, NFOL, MFOL, MOL) based on the DEGs (Supplemental Table 8). MFOL and MOL subtypes are only present in the P21 dataset because these more mature OL populations emerge after P7. Annotated ExN and OL subtypes were subsequently used to predict cell type-specific cell-cell interactions.

#### Predicting MOL subtype labels using random forest classifier

To assign MOL subtype labels from a previously published dataset^13^ to MOLs in our layer-dissected scRNA-seq data, we used the Marques et al. MOL categories to train a random forest (RF) classifier. As the Marques et al. study spans multiple brain regions, we first extracted all MOLs of cortical origin to create a reference dataset. One of the mature oligodendrocyte populations in the reference dataset (MOL2, a white-matter associated MOL) was removed from the comparison because of its very low (9 cells) representation in the cortical samples in the Marques et al. dataset. As the number of cells in each Marques et al. MOL population varied widely (Supplementary Table 5), the reference data was then downsampled to have an equal number of cells per MOL subtype; this retained 165 out of an initial 429 cells. Variable genes from the two datasets were combined and used to train the model. Training was performed with the tuneRF function in the randomForest R package (Liaw and Wiener 2002) v4.6-14, with doBest = T. For validation, 20% of the cells were reserved as a cross-validation set, and compared to a null model prepared by shuffling the cell type labels on the training data set. The trained classifier was then applied to predict MOL subtypes.

To compare between RF prediction and manually-labeled MOL subtypes, we applied Spearman correlation using the top 200 DE genes of manually identified MOL subclusters. Only the genes with adjusted p-value of < 0.05 and effect size of > 0.5 were used. Correlation is visualized using corrplot R package v0.84 (Supplemental Table 9).

#### Co-embedding OLs from different datasets using Azimuth

To visualize our scRNA-seq dataset of cortical layer dissociated OLs together with cortical OLs from the Marques et al. dataset, we applied the Azimuth algorithm via Seurat (R package version 4.0.0). UMAP was first computed on our data using the first 20 Harmony reduced dimensions with return.model=T. Between the two datasets, a set of anchors (mutual nearest neighbor cells) was identified using FindTransferAnchors function with reduction=‘pcaproject’ and reference.reduction=‘spca’, dims=1:50. IntegrateEmbeddings function followed by ProjectUMAP function were used to integrate the embeddings using the identified anchor set and to obtain projection onto the reference UMAP for each query cell.

### Slide-seq data analysis

We used a previously-published *in situ* transcriptomics (Slide-seqv2) dataset of P56 mouse somatosensory cortex in coronal section^22^. The region corresponding to the cortex was isolated from the whole puck.

Leveraging the distinct gene expression profiles and spatial connectedness of different cortical layers, beads were grouped into spatial regions (L1/Pia, L2-3, L4, L5, L6, L6b and WM) using “find_regions” function from TACCO^62^ (v0.2.2 using python v3.9.13 with annotation_key=None, position_weight=1.5, and resolution=0.7). To deconvolve cell types present in each bead, we applied “annotate” function from TACCO (with max_annotation=5 and min_log2foldchange=1).

There were two rounds of cell type deconvolution. First, we used the reference dataset from Allen Brain^26^ and subset it to include somatosensory only, to identify OL-containing beads along with other cell types. Then, to label distinct OL types and MOL subtypes, we used gene sets identified from our scRNA-seq data of OLs. This second round of deconvolution only affected beads that had some contribution of OL from the first round, by replacing OL contribution with the further-deconvoluted OL and MOL subtypes. Lastly, the density of each cell type as a function of distance from the pia was computed using “annotation_coordinate” function (with max_distance=2000 and delta_distsance=100) and plotted using a modified version of “plot_annotation_coordinate” function (available on Github).

Cells in the Slide-seq dataset were annotated using the curated marker genes for each cell type of interest (marker genes used are listed in Supplemental Table 4). Modules are plotted in their spatial location using the SpatialFeaturePlot function of Seurat v4.0.0 in R v4.0.3 with min.cutoff=’q10’.

### CellPhoneDB

Differential ligand-receptor (L-R) expression and interaction predictions were performed between PN subtypes and OLs. We filtered P7 and P21 scRNA-seq datasets generated in the laboratory and the P56 single cell dataset from the Allen Institute^26^ to systematically identify L-R pairs. We applied CellphoneDB version 2.0^27^ to obtain the interactomes at the three time points. CellphoneDB was executed by invoking the statistical_analysis method with the following parameters: --subsampling --subsampling-log=true –threads=8. Since CellphoneDB generates raw *P*-values as output, the Benjamini and Hochberg method was applied to correct for multiple testing. Output interactions were filtered using an adjusted *P* threshold of 0.05. Predicted interactions are inferred based on the mean expression of the cognate ligands and receptors from RNA-seq data of each PN sub-type and OL state. This approach captures high-confidence interaction predictions; however, weakly-expressed transcripts or drop-out expression values may result in missed predictions. Because CellPhoneDB is built using human references, we converted mouse gene symbols to human Ensembl IDs using the getLDS function from biomaRt package version 2.40.5. Human gene names in the returned predictions were then converted back to mouse gene names.

### *In utero* overexpression screen

We selected candidates predicted to promote myelination, namely genes that were more highly expressed in deep layer neurons. We chose to focus on candidates that may promote myelination, because our ligand and receptor interaction analysis is more likely to identify attractive interactions, and pro-myelination factors are also clinically more relevant for future studies.

Vectors containing the sequence of the target genes selected from the L-R interaction analysis were purchased from Horizon discovery. Target genes were subcloned by PCR-based cloning (NEBuilder HiFi DNA cloning, E5520S, New England BioLabs) into a mammalian expression vector (PCM153) under the control of the EF1a promoter, which allows robust and long-term expression of the transgene. The expression vector containing the target gene of interest was co-electroporated by *in-utero* electroporation with the pCAG-EGFP vector to obtain a strong GFP fluorescent signal that allows clear identification of the electroporated neurons (Figure 5A). For the negative control, the empty expression vector was co-electroporated with pCAG-EGFP.

*In utero* electroporation (IUE) was performed as previously described^63–65^. In brief, purified DNA (2 µg/µl) mixed with 0.005% fast green in sterile PBS was injected *in utero* into the lateral ventricle of CD1 embryos. Five pulses of 40V and 50ms were delivered at 1s intervals with a CYU21EDIT square wave electroporator and 1cm diameter platinum electrodes (Napa Gene).

Electroporated progeny were analyzed at different postnatal stages: P14 (which corresponds to the initial phase of myelination in deep layers), and P21 (when the mouse cortex is actively being myelinated).

### Fluorescence immunolabelling

Immunohistochemistry was performed using P14 and P21 mice. Animals were anesthetized with tribromoethanol (Avertin) and transcardially perfused with 0.1 M PBS (phosphate buffered saline, pH 7.4) followed by ice-cold 4% paraformaldehyde (Electron Microscopy Sciences, 15710), as previously described^56^. Cortical tissue was then post-fixed overnight in 4% paraformaldehyde, followed by 3X 10-minutes washes in 0.1 M PBS. Serial coronal sections (40 um thick) were cut using a Leica microtome (VT1000 S), collected in PBS and stored at 4°C. For most comparative analysis, sectioned matched, primary somatosensory cortex (S1 cortex) tissues were used for immunohistochemistry. Free-floating sections were blocked for 1-2 hour at room temperature in blocking buffer (PBS with 0.3-0.5% BSA, 0.3% Triton X-100, and 2-8% donkey serum), and then incubated overnight at 4°C in blocking buffer with the following primary antibodies: rat anti-MBP (1:100, MAB386, Millipore), rabbit anti-Cux1 (1:100, CDP M-222, Santa Cruz Biotechnology), rat anti-Ctip2 (1:100, ab18465, Abcam), mouse anti-APC (1:100, OP80, Millipore), goat anti-PDGFRa (1:100, Novus Biologicals, AF1062), rabbit anti-GFP (1:500, A11122, Invitrogen), mouse anti-MBP (BioLegend, previously Covance #SMI-99P). Secondary antibody labeling was performed in blocking buffer at room temperature for 2 hours as follows: donkey anti-rat IgG (H+L) Alexa Fluor 647 preabsorbed (1:500-800, ab150155, Abcam), donkey anti-mouse IgG (H+L) Alexa Fluor 546 (1:200, A10036, Thermo Fisher Scientific), donkey anti-rabbit IgG (H+L) Alexa Fluor 488 (1:200, a21206, Thermo Fisher Scientific), donkey anti-goat IgG (H+L) Alexa Fluor 546 (1:200, A11056, Thermo Fisher Scientific). Sections were mounted using Fluoromount-G (00-4958-02, Thermo Fisher Scientific).

### Imaging

Wide-field fluorescence images were acquired using a Zeiss Axio Imager 2 upright microscope. Tile scan images spanning the entire brain slice were captured using a 10x air objective and stitched using Zeiss Zen Blue Software.

### Image Analysis

#### Reelin mutant and control mice

For every cortical section, we divided the cortex from the pia to the white matter into 10 horizontal bins, made the plot profile, and calculated the average pixel value of fluorescence for MPB, CTIP2, and CUX1 in each bin using Fiji^66^. The values for each bin were averaged for 9 sections (3 sections each from 3 individual animals from the S1 cortex) for each of the WT and *Reeler^-/-^* to obtain an average plot profile.

Pearson’s correlation coefficient was computed for the average plot profile for MBP, CTIP2 and CUX1.

#### *In vivo* overexpression screen

For every tissue section, electroporated and contralateral regions of the S1 cortex, from pia to corpus callosum, were analyzed. Cortical thickness was divided in two equal-height bins, termed Upper layers (UL) and Deep layers (DL). To quantify MBP protein expression levels, we calculated the average pixel value per bin using Fiji (v 2.9.0/1.53t). To quantify APC density, images were thresholded, and masked APC^+^ cell bodies automatically counted using the ‘analyze particles’ function of Fiji. APC density per mm^2^ was calculated per bin.

The significance of the differences in the fluorescence intensities of MBP and APC density between IUE and contralateral was computed using linear models. For each candidate gene tested, linear models were fit on the data, modeling biological replicate (animal) and section as fixed effects, and with or without the electroporation as another fixed effect. Two fitted models were compared using the ANOVA function to evaluate the use of an additional predictor.

## ACKNOWLEDGMENTS

We thank Sung Min Yang, Jeffrey A. Stogsdill, Zachary Trayes-Gibson, Lila Luna Lyons, Sarah Sime, and former and present members of the Arlotta laboratory for insightful discussions and experimental help; Douglas Richardson for helpful advice on imaging; and Nathan Curry and Zachary Niziolek for excellent technical support with sorting. This work was supported by grants from the Broad Institute of MIT and Harvard and the National Institute of Health (R01NS128117 and R01NS103758) to P.A., the Stanley Center for Psychiatric Research to P.A. and J.Z.L., and the Charles A. King Trust postdoctoral research fellowship to V.J.

## AUTHOR CONTRIBUTIONS

P.A. and V.J. conceived the experiments. V.J., N.D.I., A.S., W.Y. and X.J. performed the experiments, with help from P.O.C. and C.A.. K.K., S.S., D.D.B. and X.J. analyzed the transcriptomic data, with guidance from J.Z.L.. P.A., J.B., V.J. and N.D.I. wrote the manuscript with contributions from all authors.

