## Supplemental Information for "Neuronal-class specific molecular cues drive differential myelination in the neocortex"

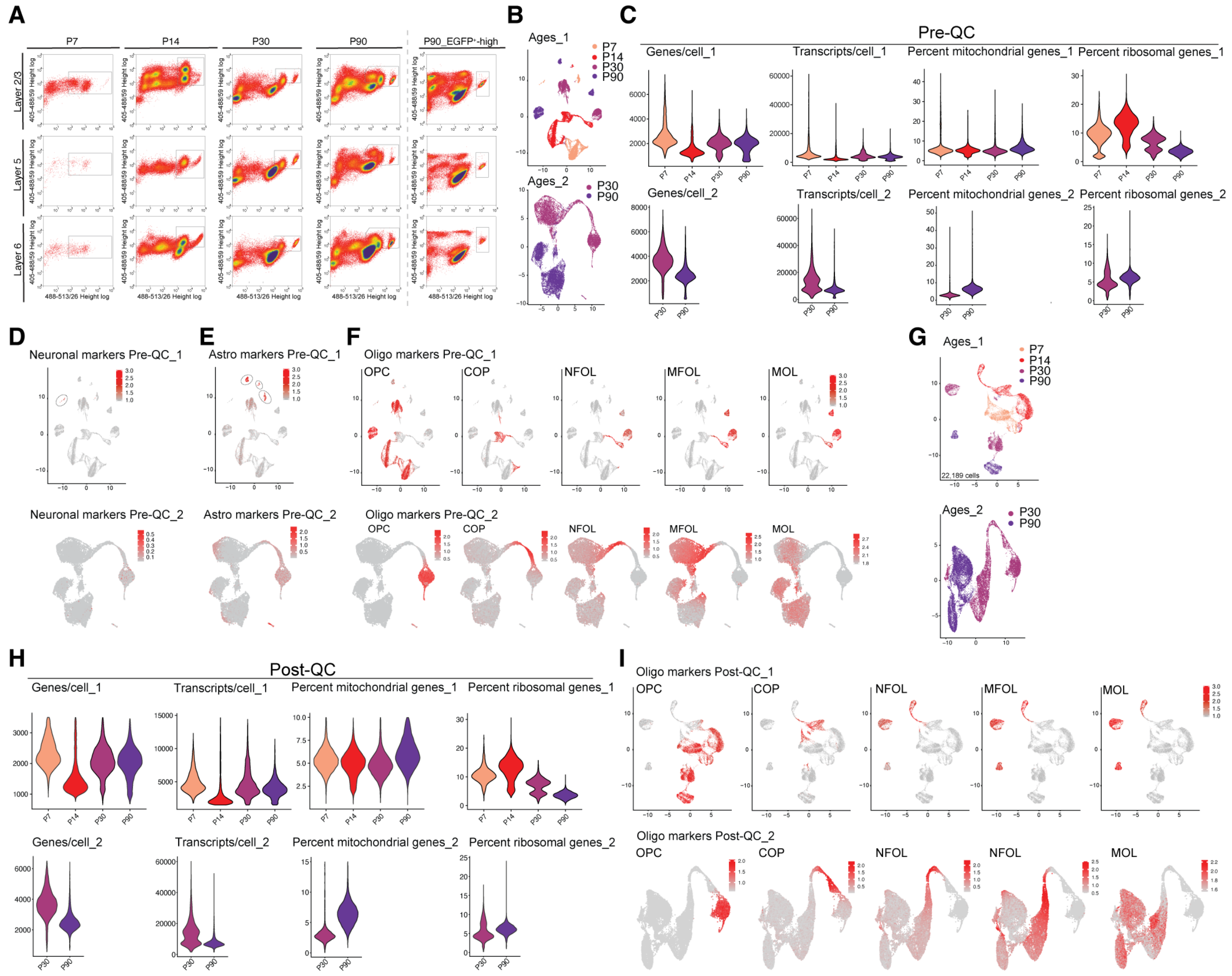


**Figure S1. Cell sorting and quality control analysis of scRNA-seq of oligodendrocytes from *Plp1-EGFP*$^{+/-}$ micro-dissected cortical layers.**

(**A**) FACS plots of EGFP$^{+}$ oligodendrocyte sorting from P7, P14, P30 and P90 *Plp1-EGFP*$^{+/-}$ mice. S1 cortices were micro-dissected and cells from L2/3 (top), L5 (middle), and L6 (bottom) were sorted separately. FACS gating for live cells (Hoechst$^{+}$) and EGFP$^{+}$ cells is shown by boxes. For a separate experimental sample at P90, the population with the highest EGFP signal was sorted (P90_EGFP$^{+}$-high). Only live, EGFP$^{+}$ cells were used for scRNA-seq experiments, to enrich oligodendrocyte-lineage cells.

(**B**) UMAP of *Plp1-EGFP*$^{+}$ scRNA-seq datasets for each replicate, color-coded by age.

(**C**) Violin plots of initial quality control metrics of the P7, P14, P30, and P90 *Plp1-EGFP*$^{+}$ scRNA-seq datasets for each replicate before removal of doublets, low-quality cells, and non-OL cells (Pre-QC). Genes/cell, Transcripts/cell, percent of mitochondrial gene expression/cell, and percent of ribosomal gene expression/cell are plotted. Data are split and color-coded by age.

(**D**) Expression feature plots for the neuronal gene module (see Supplemental Table 7) for all ages combined in *Plp1-EGFP*$^{+}$ scRNA-seq datasets for each replicate. This identified one neuronal cluster (circled), which was removed for the final analysis.

(**E**) Expression feature plots for the astrocyte gene module (see Supplemental Table 7) for all ages combined in *Plp1-EGFP*$^{+}$ scRNA-seq datasets for each replicate. This identified three astrocytic cell clusters (circled), which were removed for the final analysis.

(**F**) Expression feature plots for OL gene modules (see Supplemental Table 7) for all ages combined in *Plp1-EGFP*$^{+}$ scRNA-seq datasets Pre-QC for each replicate. Gene modules represent stages of OL maturation: OPC (OL progenitor cell), COP (committed OL progenitor), NFOL (newly formed OL), MFOL (myelin forming OL), and MOL (myelinating OL).

(**G**) UMAP of *Plp1-EGFP*$^{+}$ scRNA-seq clusters for each replicate after removal of doublets, low-quality cells, and non-OL cells (Post-QC), color-coded by age.

(**H**) Violin plots of quality control metrics of the P7, P14, P30, and P90 *Plp1-EGFP*$^{+}$ scRNA-seq datasets for each replicate after removal of doublets, low-quality cells, and non-OL cells (Post-QC). Genes/cell, Transcripts/cell, percent of mitochondrial gene expression/cell, and percent of ribosomal gene expression/cell are plotted. Data are split and color-coded by age.

(**I**) Expression feature plots for oligodendrocyte gene module expression for all ages combined in *Plp1-EGFP*$^{+}$ scRNA-seq datasets Post-QC for each replicate. Gene modules represent stages of oligodendrocyte maturation: OPC (oligodendrocyte progenitor cell), COP (committed oligodendrocyte progenitor), NFOL (newly formed oligodendrocytes), MFOL (myelin forming oligodendrocytes), and MOL (myelinating oligodendrocytes).

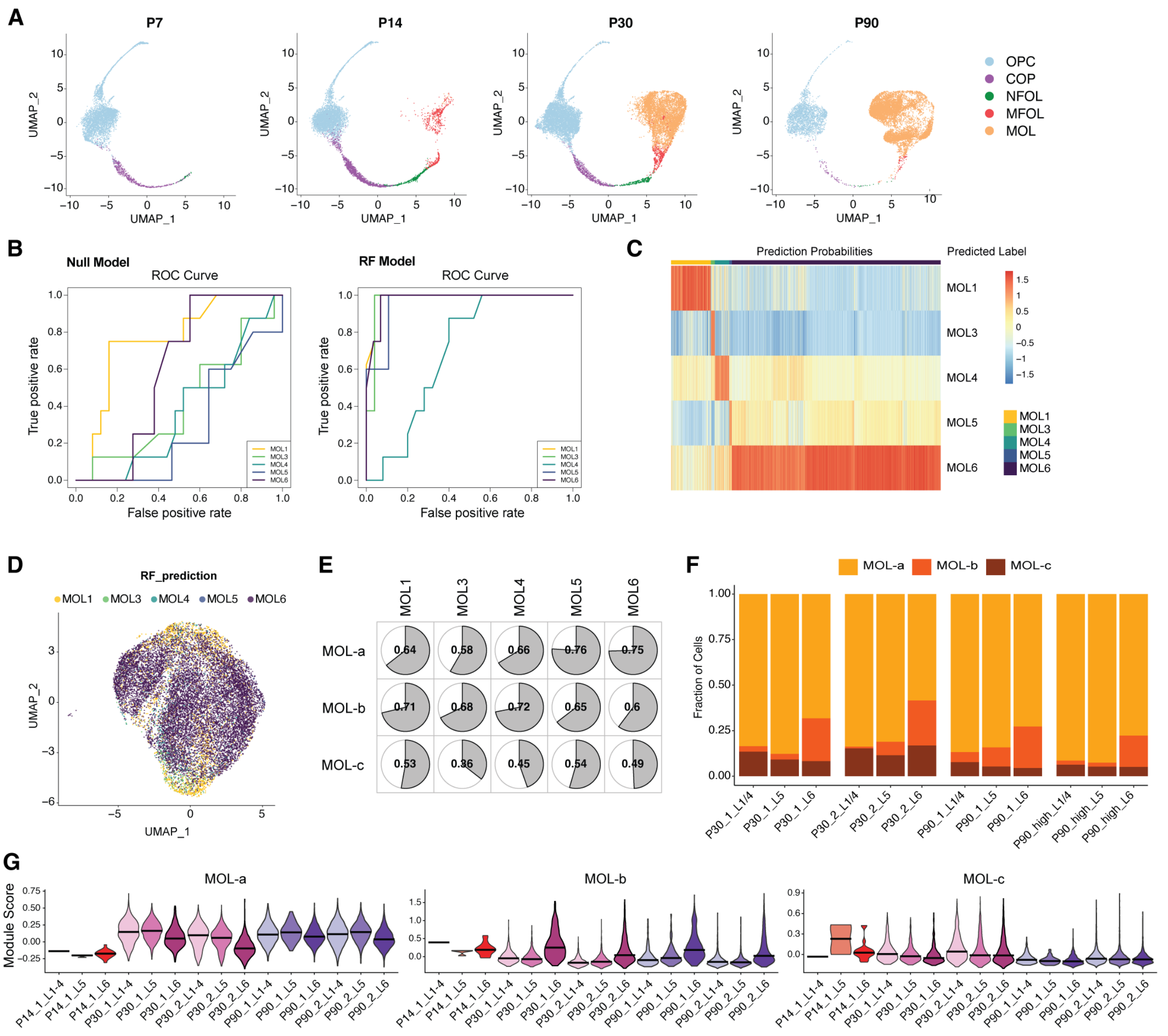

**Figure S2. MOL states across time and cortical layers.**

(**A**) UMAP of OL-lineage cells, color-coded by age (both experiments combined).

(**B**) Receiver-operating-characteristic curves comparing the performance of the trained random forest classifier to that of a null model (see STAR Methods).

(**C**) Heatmap showing the prediction confidence of the RF model we used to re-assign our MOL subtypes to those defined by Marques et al., 2016.

(**D**) UMAP showing the re-assigned MOL subtypes (color-coded) according to the RF prediction result.

(**E**) Pearson correlation coefficient for correspondence between each of the MOL states identified in this study (MOL a-c) with the MOL states identified in Marques et al., 2016 (MOL1-6).

(**F**) Bar plot of P30 and P90 MOL substate proportions for each micro-dissected layer. Color-coding indicates the MOL sub-types. Data are normalized to the total cell count in each micro-dissected layer.

(**G**) Violin plots of MOL-a, MOL-b and MOL-c module score in all cell types in the scRNA-seq data.

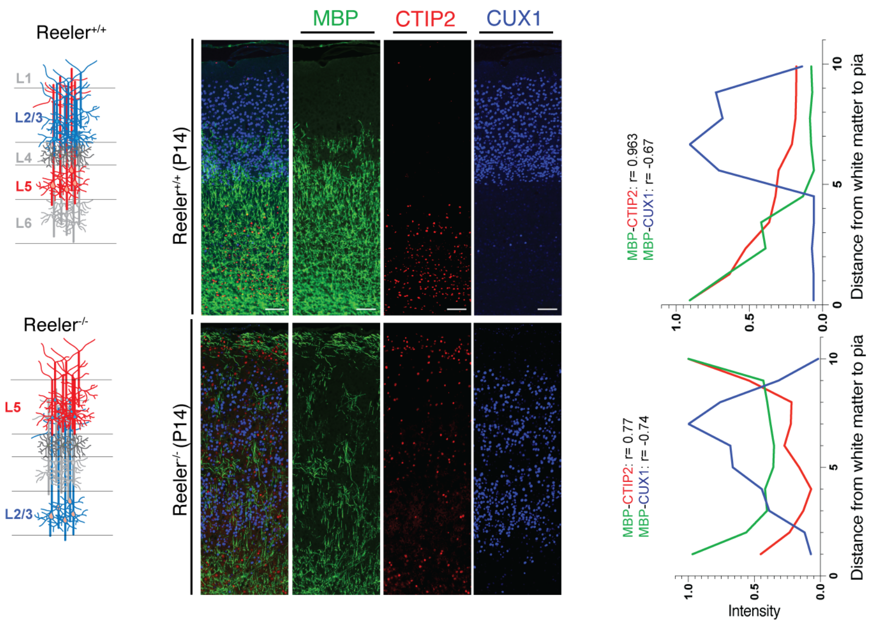


**Figure S3. Projection neuron subtype and not location determine myelination.**

Left, schematic representation of varying PN subtype distribution in the WT (*Reln*$^{+/+}$) and Reln$^{-/-}$ mouse cortical plate. Center, representative images of S1 cortical region of P14 WT (top) and *Reln*$^{-/-}$ (bottom) mice immunolabelled for MBP (green), CTIP2 (red) and CUX1 (blue) (scale bar: 100µm) (n= 3mice, 3 sections). Right, quantification of MBP, CTIP2 and CUX1 intensities along the cortex (r = Pearson correlation of the two indicated intensities along the depths of the cortex).

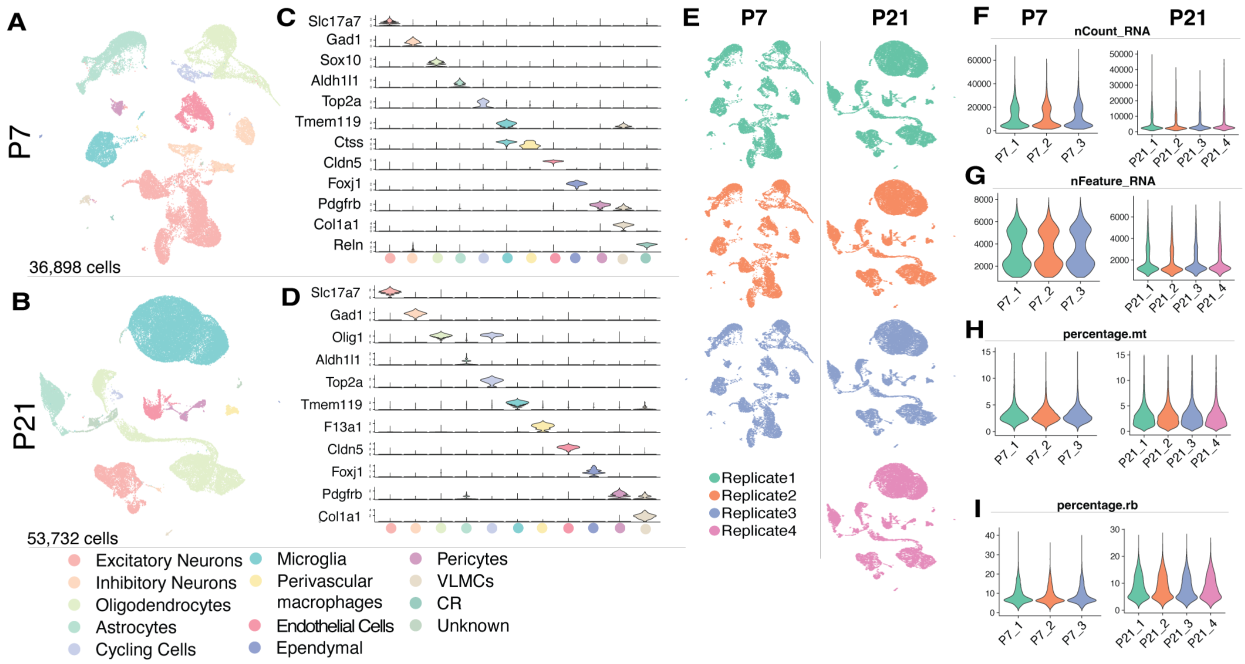


**Figure S4. Analysis of scRNA-sequencing datasets from mouse somatosensory cortex at P7 and P21.**

(**A-B**), UMAP of all-lineage cells isolated from cortex, color-coded by subtype at P7 and P21.

(**C-D**) Violin plot of expression of key marker genes across different cell populations.

(**E**) UMAP of replicates at P7 (left) and P21 (right).

(**F-I**) Violin plots of initial quality control metrics of the P7 (left) and P21 (right) scRNA-seq datasets after removal of doublets and low-quality cells, and keeping only OL or PN lineages. Genes/cell, Transcripts/cell, percent of mitochondrial gene expression/cell, and percent of ribosomal gene expression/cell are plotted. Data are split and color-coded by replicate.

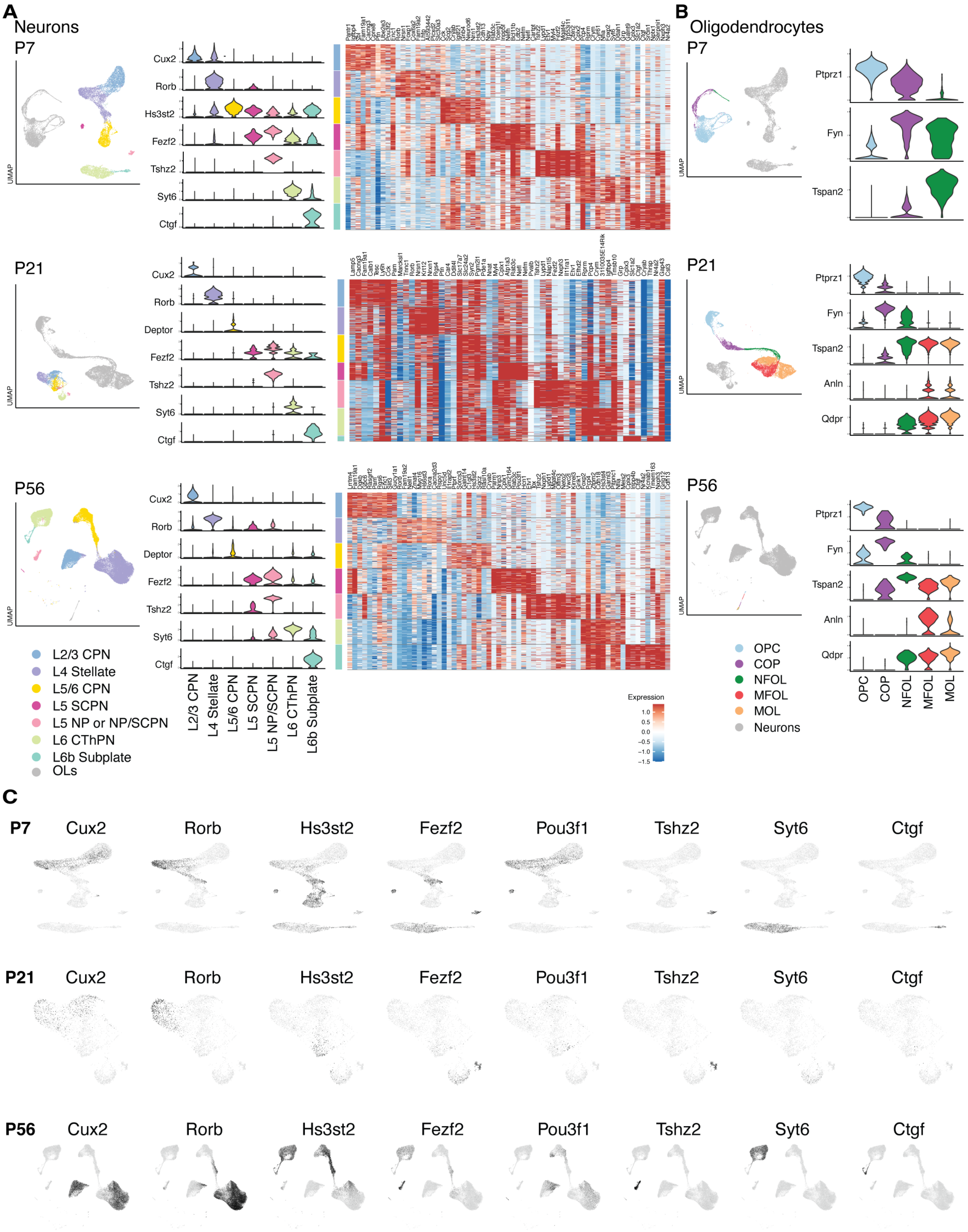
A
Neurons
P7
UMAP
Cux2
Rorb
Hs3st2
Fezf2
Tshz2
Syt6
Ctgf
P21
Deptor
P56
L2/3 CPN
L4 Stellate
L5/6 CPN
L5 SCPN
L5 NP or NP/SCPN
L6 CThPN
L6b Subplate
OLs
L5 NP/SCPN
Expression
B
Oligodendrocytes
Ptprz1
Fyn
Tspan2
Anln
Qdpr
OPC
COP
NFOL
MFOL
MOL
Neurons
C
Pou3f1

**Figure S5. Analysis of neuronal and oligodendrocyte populations in scRNA-sequencing data from mouse somatosensory cortex.**

(**A**) UMAP of PN-lineage cells, color-coded by subtype at P7 and P21 (this manuscript) and P56 (from Allen Brain) (left). Violin plots of the expression of key marker genes across different PN populations (middle) and heatmaps of the top 10 DEGs (by MAST) for each PN population (right).

(**B**) UMAP of OL-lineage cells, color-coded by subtype at P7 and P21 (this manuscript) and P56 (from Allen Brain) (left). Violin plots of expression of key marker genes across different OL populations (right).

(**C**) Expression feature plots for marker genes of neuronal sub-types at P7, P21, and P56.

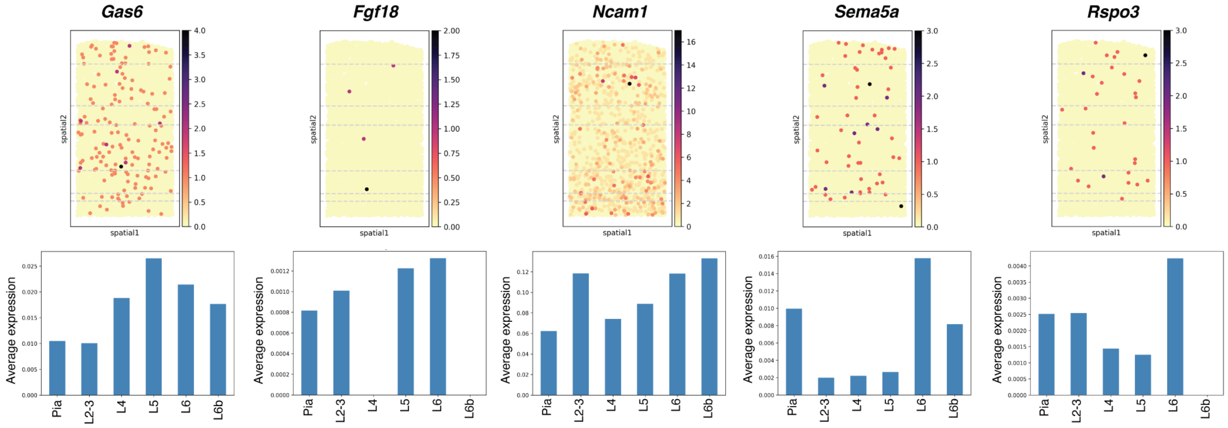


**Figure S6. Visualizing candidate gene expression across cortical layers by in situ transcriptomics.**

Gene counts of each candidate gene plotted onto the Slide-seq data (P56 mouse cortical section) (top; dashed lines indicate cortical layers), with average gene expression for each candidate gene in each layer (bottom).

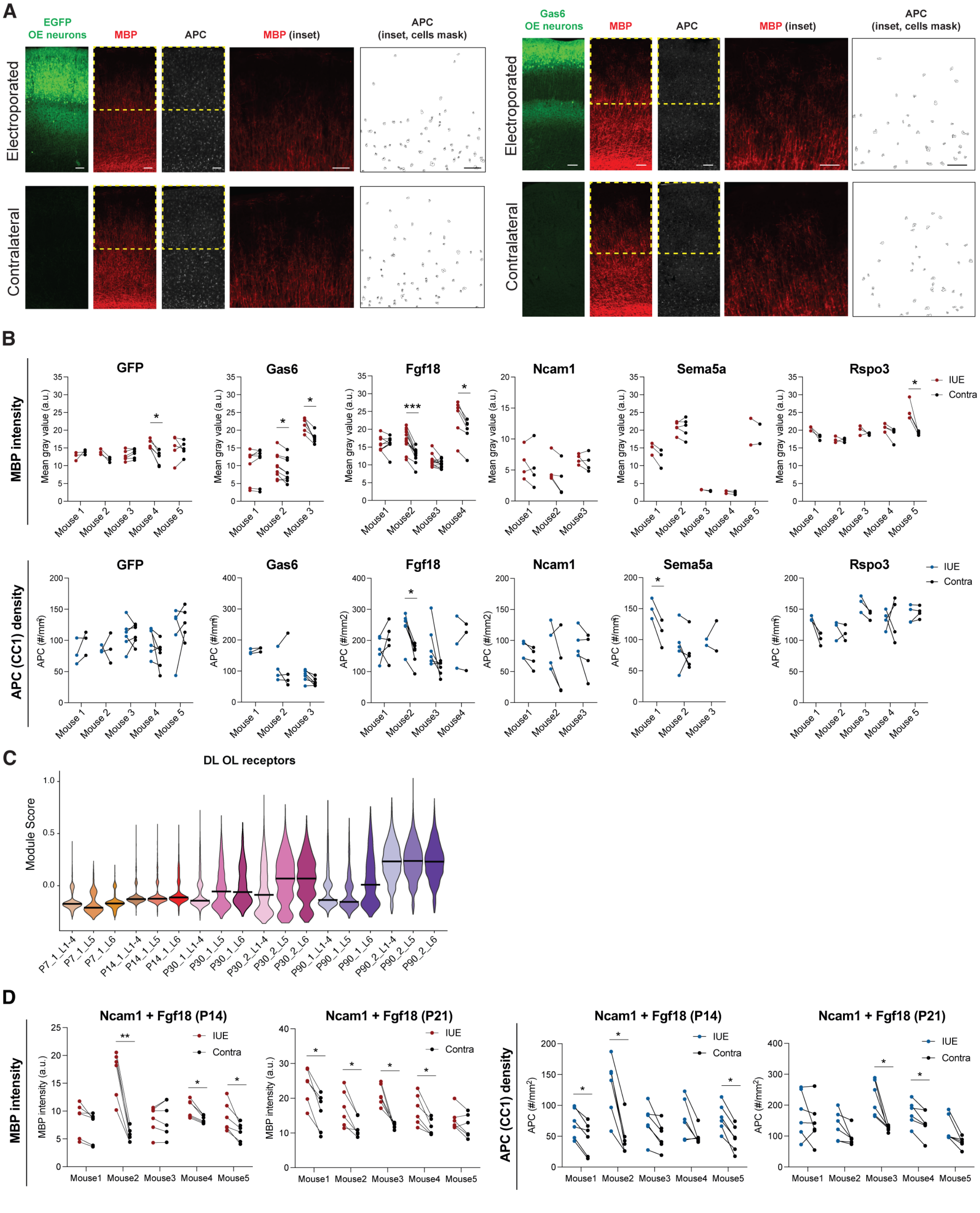


**Figure S7. Modulation of myelination after overexpression of candidate genes**

(**A**) Representative images of IUE and contralateral S1 cortical regions at P21 analyzed for MBP intensity (red) and APC$^{+}$ cell density (blue), for negative control (*EGFP*) and positive control

(*Gas6* over-expression). Sections were immunolabelled for MBP in red and APC (CC1, mature oligodendrocyte marker) in blue. Electroporated cells express EGFP. Analyzed electroporated and contralateral regions are highlighted with a yellow box (scale bar: 500µm). Inset: magnification of upper cortical layers, showing MBP intensity and APC count (scale bar: 100µm).

(**B**) Quantification of myelin basic protein (MBP) intensity (top) and $APC^+$ (CC1, mature oligodendrocyte marker) cell density in upper layers (bottom) of electroporated (IUE, red for MBP and blue for APC) and contralateral (contra, black) cortical regions at P21 after *in utero* electroporation (IUE) of candidate mediators of myelination. Each data point represents one brain section. (* p<0.05; ** p<0.01; *** p<0.001).

Number of sections and p-values from linear models, MBP:

*EGFP:* mouse1=3 sections, mouse2=3 sections, mouse3=6 sections, mouse4=5 sections, mouse5=6 sections (p-values from linear models, mouse1 p=0.483, mouse2 p=0.083, mouse3 p=0.100, mouse4 p=0.0115, mouse5 p=0.902)

*Fgf18:* mouse1=9 sections, mouse2=12 sections, mouse3=12 sections, mouse4=6 sections (p-values from linear models, mouse1 p=0.957; mouse2 p=0.00016, mouse3 p=0.098, mouse4 p=0.011).

*Ncam1:* mouse1=4 sections, mouse2=4 sections, mouse3=4 sections (p-values from linear models, mouse1 p=0.687, mouse2 p=0.099, mouse3 p=0.492).

*Gas6:* mouse1=7 sections, mouse2=9 sections, mouse3=6 sections (p-values from linear models, mouse1 p=0.469, mouse2= 0.021, mouse3 p=0.021).

*Sema5a:* mouse1=3 sections, mouse2=5 sections, mouse3=2 sections, mouse4=3 sections, mouse5=2 sections (p-values from linear models, mouse1 p=0.089, mouse2 p=0.912, mouse3 p=0.367, mouse4 p=0.231, mouse5 p=0.755).

*Rspo3:* mouse1=3 sections, mouse2=3 sections, mouse3=3 sections, mouse4=4 sections, mouse5=4 sections (p-values from linear models, mouse1 p=0.099, mouse2 p=0.875, mouse3 p=0.492, mouse4 p=0.098, mouse5 p=0.045).

Number of sections and p-values from linear models, APC:

*EGFP:* mouse1=3 sections, mouse2=3 sections, mouse3=6 sections, mouse4=5 sections, mouse5=6 sections (p-values from linear models, mouse1 p=0.413, mouse2 p=0.979, mouse3 p=0.979, mouse 4 p=0.261, mouse5 p=0.636).

*Fgf18:* mouse1=5 sections, mouse2=6 sections, mouse3=6 sections, mouse4=3 sections (mouse1 p=0.536, mouse2 p=0.018, mouse3 p=0.131, mouse4 p=0.979).

*Ncam1:* mouse1=4 sections, mouse2=4 sections, mouse3=4 sections (p-values from linear models, mouse1 p=0.125, mouse2 p=0.300; mouse3 p=0.313).

*Gas6* mouse1=3 sections, mouse2=4 sections, mouse3=6 sections (p-values from linear models, mouse1 p=0.309, mouse2 p=0.979, mouse3 p=0.076).

*Sema5a:* mouse1=3 sections, mouse2=5 sections, mouse5=2 sections (p-values from linear models, mouse1 p=0.046, mouse2 p=0.615, mouse3 p=0.809).

*Rspo3:* mouse1=3 sections, mouse2=3 sections, mouse3=3 sections, mouse4=4 sections, mouse5=4 sections (p-values from linear models, mouse1 p=0.108, mouse2 p=0.801, mouse3 p=0.125, mouse4 p=0.995, mouse5 p=0.544

(**C**) Violin plot of gene module score for corresponding DL OL receptor genes (*Lgr4, Fgfr1,* and *Fgfr2*) by age and layer for the oligodendrocyte scRNA-seq data. The black line represents the median.

(**D**) Quantification at P14 and P21 of myelin basic protein (MBP) intensity (left) and $APC^{+}$ cell density (right) in upper layers of electroporated (IUE, red for MBP and blue for APC) and contralateral (contra, black) cortical regions after IUE. Each data point represents one brain section (P14: n=5 mice, N=30 sections total, P21: n=5 mice, N=30 sections total) (* p<0.05; ** p<0.01; *** p<0.001).

p-values from linear models:

P14:

MBP: mouse1 p=0.083; mouse2 p=0.006; mouse3 p=0.786; mouse4 p=0.021; mouse5 p=0.029.

APC: mouse1 p=0.018; mouse2 p=0.016; mouse3 p=0.072; mouse4 p=0.125; mouse5 p=0.0125.

P21:

MBP: mouse1 p=0.011; mouse2 p=0.029; mouse3 p=0.011; mouse4 p=0.021; mouse5 p=0.158.

APC: mouse1 p=0.508; mouse2 p=0.136; mouse3 p=0.046; mouse4 p=0.046; mouse5 p=0.076).
